# The SPOC domain is a phosphoserine binding module that bridges transcription machinery with co- and post-transcriptional regulators

**DOI:** 10.1101/2022.02.26.482114

**Authors:** Lisa-Marie Appel, Irina Grishkovskaya, Johannes Benedum, Vedran Franke, Anton Polyansky, Andrea Neudolt, Anna Wunder, Bojan Zagrovic, Altuna Akalin, Kristina Djinovic-Carugo, Dea Slade

## Abstract

The heptarepeats of the C-terminal domain (CTD) of RNA polymerase II (Pol II) are extensively modified throughout the transcription cycle. The CTD coordinates RNA synthesis and processing by recruiting transcription regulation factors as well as RNA capping, splicing and 3’end processing factors. The SPOC domain of PHF3 was recently identified as a new CTD reader domain specifically binding to phosphorylated serine-2 residues in adjacent CTD repeats. Here, we establish the SPOC domains of the human proteins DIDO, SHARP and RBM15 as phosphoserine binding modules that can act as CTD readers but also recognize other phosphorylated binding partners. We report the crystal structure of SHARP (SPEN) SPOC-CTD and identify the molecular determinants for its specific binding to phosphorylated serine-5. PHF3 and DIDO SPOC domains preferentially interact with the Pol II elongation complex, while RBM15 and SHARP SPOC domains engage with the m6A writer and reader proteins. Our findings establish the SPOC domain as a major interface between the transcription machinery and regulators of transcription and co-transcriptional processes.

## Introduction

The SPOC (Spen orthologue and paralogue C-terminal) domain is a 15-20 kDa protein domain found across eukaryotic species from yeast to mammals^1^. SPOC domains form a distorted β-barrel structure comprising seven β-strands and a variable number of α-helices^2–5^. The seven human SPOC containing proteins can be divided into three groups based on their domain organization: Spen family, DIDO/PHF3 and SPOCD1. SHARP (SMRT/HDAC1 associated repressor protein), RBM15 and RBM15B (RNA binding motif protein 15/15B) paralogues belong to the Spen family of proteins characterized by a series of N-terminal RRMs (RNA recognition motifs) and the C-terminal SPOC domain. DIDO (Death inducer obliterator) and PHF3 (PHD finger protein 3) share a different domain architecture comprising a PHD (Plant homeodomain), a TLD (TFIIS like domain) and the SPOC domain. SPOCD1 (SPOC domain containing protein 1) originated from a duplication of PHF3^6^ but lacks the PHD.

SPOC containing proteins are generally associated with transcription regulation, differentiation and development^5,7–10^. The SPOC domain of SHARP is crucial for its repressive function in the Notch signaling pathway. It recruits corepressor complexes SMRT/NCoR-HDAC1 by binding to phosphorylated serine within the conserved LSD motif at the SMRT/NCoR C-terminus^3, 11^. More recently, SHARP SPOC has been implicated in Xist lncRNA-mediated silencing during X-chromosome inactivation but the mechanism remains elusive^12^. In comparison to SHARP SPOC, the SPOC domains of the other two Spen family members, RBM15 and RBM15B, have a weaker effect in repressing transcription when tethered to a promoter through Gal4-DBD^13^. Instead, RBM15 and RBM15B are involved in post-transcriptional regulation, mainly by influencing alternative splicing, m6A (N^6^-methyladenosine) RNA modification, and nuclear export^10, 14–16^. Little is known about the function of their SPOC domains. RBM15 SPOC was shown to bind to the unstructured LPDSD motif of the histone H3K4me3 methyltransferase^17^. Additionally, RBM15 and RBM15B SPOC domains were shown to bind to the Epstein-Barr virus early protein EB2, which promotes nuclear export of viral mRNAs^13^.

We recently showed that PHF3 SPOC specifically binds the C-terminal domain (CTD) of the RNA polymerase II (Pol II) subunit RPB1 phosphorylated on serine-2 in tandem repeats^5^. The disordered CTD comprises up to 52 heptarepeats of the sequence YSPTSPS and is differentially modified throughout the transcription cycle^18, 19^. Phosphorylation of serine-5 is a mark of early stages of transcription while productive elongation is linked with serine-2 phosphorylation^18^. Different phosphomarks are recognized by CTD reader domains, ensuring the timely recruitment of transcription regulators and RNA processing factors to the transcription machinery^19–21^. The SPOC domain is critical for PHF3-mediated regulation of transcription and mRNA stability of neuronal genes^5^. The PHF3 paralogue DIDO regulates splicing and is critical for stem cell self-renewal and differentiation^22–25^, while SPOCD1 is required for silencing of transposable elements through pi-RNA mediated methylation^6^. However, the function of DIDO and SPOCD1 SPOC domains has remained unclear.

SHARP and PHF3 SPOC have been established as phosphoserine binding domains raising the question as to whether other SPOC domains also bind phosphorylated serine and how they contribute to protein interaction networks. Here we show that the SPOC domains of DIDO, SHARP and RBM15 act as CTD reader domains that bind phosphorylated serines via conserved surface patches. We report the crystal structure of SHARP in complex with serine-5-phosphorylated CTD and determine the similarities and differences between SHARP-SMRT and SHARP-CTD interaction on the structural level. We further applied mass spectrometry to identify SPOC-dependent interactors of PHF3, DIDO, SHARP and RBM15. Our findings establish Pol II elongation machinery as the focal point for PHF3 and DIDO SPOC interactions, while m6A writer and reader proteins are major targets for RBM15 and SHARP SPOC domains. Collectively, our results suggest that SPOC is a versatile phosphoserine binding module that spans the transcription machinery, co- and post-transcriptional regulators.

## Results

### Conserved surfaces on SPOC mediate binding to phosphorylated serine

PHF3 SPOC has been established as a CTD reader domain^5^, but it has remained unknown whether CTD binding is unique to PHF3 or other SPOC domains possess the same ability. SPOC domains show an overall low level of sequence conservation, however, some key residues are conserved (Fig. 1a). These include an arginine residue (red asterisk in Fig. 1a) that is conserved in all human SPOC domains except SPOCD1 and is critical for the electrostatic anchoring of phosphorylated serine in CTD by PHF3 SPOC and in SMRT/NCoR by SHARP SPOC (R1248 in PHF3, R3552 in SHARP, Fig. 1b,c, 3e)^3, 5, 11^. This arginine residue is part of a positively charged patch on the surface of SPOC. PHF3 SPOC has two such patches (Fig. 1b) while SHARP SPOC has one basic patch (Fig. 1c), which contain additional conserved lysine and arginine residues (Fig. 1a, conserved residues marked with red squares). Although experimental structural information on DIDO SPOC is lacking, AlphaFold2 structure prediction^26, 27^ shows that the conserved residues cluster in patches on the domain surface similar to PHF3 SPOC (Fig. 1d) suggesting that these domains may have similar phosphoserine binding properties. We solved the crystal structure of RBM15 SPOC (to 1.45 Å resolution), which has a distorted β-barrel fold comparable to previously structurally characterized SPOC domains (Fig. 1e, Supplementary Fig. 1, Supplementary Table 1). The conserved lysine and arginine residues cluster to a basic surface patch similar to that of SHARP SPOC (Fig. 1e, Supplementary Fig. 1b). Although phylogenetic analysis indicates that SPOCD1 originated from a duplication of PHF3^6^, only two out of four basic residues from PHF3 are also conserved in SPOCD1 (Fig. 1a), suggesting that SPOCD1 SPOC has lost the phosphoserine binding ability. Indeed, the structure predicted by AlphaFold2 reveals a much weaker positive charge in the surface patches of SPOCD1 SPOC (Fig. 1f). Furthermore, the distance between the patches is 35.550 Å, which is considerably further apart than the patches of PHF3 or DIDO SPOC (24.269 Å and 20.592 Å, respectively) and makes it highly unlikely that SPOCD1 SPOC could accommodate CTD phosphorylations in adjacent repeats.

**Fig. 1:**
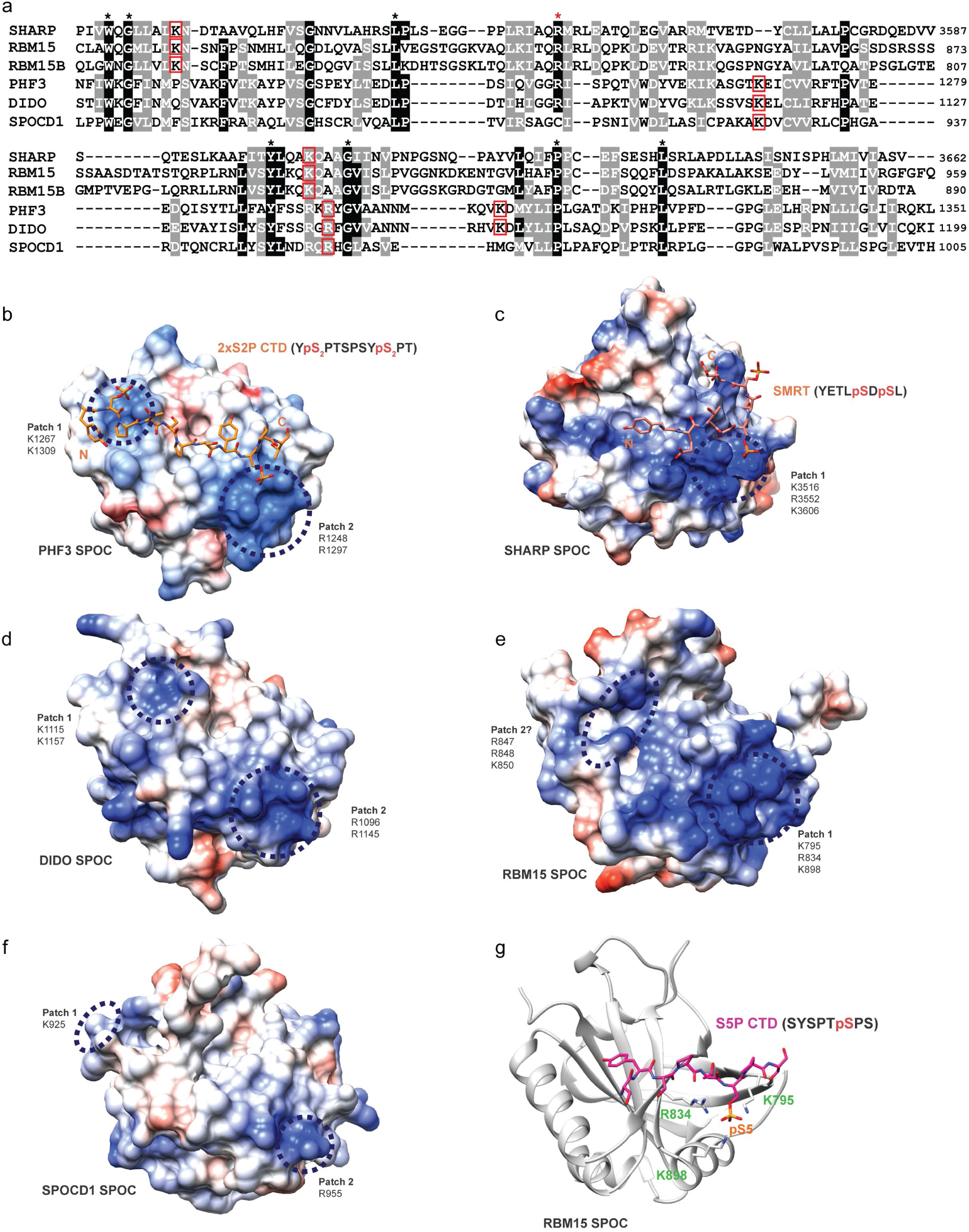
Conserved surfaces on SPOC mediate phosphoserine binding. **a** Multiple sequence alignment of human SPOC domains based on PROMALSD3 using SPOC structures from human SHARP (2RT5), human PHF3 (6Q2V) and sequences from human RBM15, RBM15B, DIDO and SPOCD1. Red squares indicate conserved residues that constitute basic patches on the surface of SPOC. Red asterisk indicates an arginine residue that is conserved in all human SPOC domains except SPOCD1. **b** Crystal structure of PHF3 SPOC in complex with 2xS2P CTD peptide (6IC8). Conserved basic patches that mediate binding to phosphorylated CTD residues are indicated with blue circles. The distance between the patches is 24.296 Å. **c** NMR solution structure of SHARP SPOC in complex with phosphorylated SMRT peptide (2RT5). The blue circle indicated the conserved basic patch that coordinates binding to SMRT pS2522. **d** AlphaFold2 structural model DIDO SPOC (Q9BTC0). Blue circles indicate conserved surface patches. The distance between the patches is 20.592 Å. **e** Crystal structure of RBM15 SPOC (7Z27). Blue circles indicate the conserved basic surface patch (patch 1) and a potential second patch (patch 2). The distance between the patches is 20.891 Å. **f** AlphaFold2 structural model of SPOCD1 SPOC (Q6ZMY3). The surface patches indicated by blue circles are only partially conserved and display a less pronounced positive charge. The distance between the patches is 35.550 Å. **g** Structural model of the interaction between RBM15 SPOC and serine-5-phosphorylated CTD generated in PyMOL and refined using the HADDOCK2.2 webserver^28, 29^. All SPOC domains are shown in the same orientation and at the same scale. Electrostatic surface potential in **b-f** was calculated using the Coulombic Surface Coloring tool in UCSF Chimera^64^ and is depicted ranging from -10 (red) to +10 (blue) kcal/(mol*e). The distances in **b** and **d-f** are given as the mean distance between the terminal atoms of the amino acids making up the basic patches and were measured using the Structure Measurements – Distances tool in UCSF Chimera.

To test whether SPOC domains are universal CTD binders and determine their binding specificity, we expressed and purified the SPOC domains of the human proteins PHF3, DIDO, SHARP, RBM15 and SPOCD1 and performed fluorescence anisotropy assays to measure their binding to Atto488-labeled CTD-diheptapeptides (Fig. 2, Supplementary Fig. 2-6). To obtain a comprehensive picture of SPOC-CTD interactions, we measured binding affinities for a total of eleven peptides that were either unphosphorylated (CTD) or phosphorylated in one (1xS2P, 1xS5P, 1xS7P, 1xY1P, 1xT4P) or both repeats (2xS2P, 2xS5P, 2xS7P, 2xY1P, 2xT4P) (Supplementary Table 2). We first confirmed our previous finding that PHF3 SPOC preferentially binds to CTD phosphorylated at serine-2 in two adjacent repeats (2xS2P, Kd = 0.42 ± 0.02 µM, Fig. 2a). PHF3 did not bind to unphosphorylated or single phosphorylated CTD peptides and bound with lower affinity to other double phosphorylations (Fig. 2a,e and Supplementary Fig. 2; Kd ranging from 29 µM to 130 µM). Similarly, the SPOC domain of the PHF3 paralogue DIDO showed highest affinity for the 2xS2P peptide (Kd = 4.8 ± 0.6 µM), did not bind unphosphorylated or single phosphorylated CTD and bound other double phosphorylated peptides with low affinity (Fig. 2b,e and Supplementary Fig. 3; Kd ranging from 102 µM to 298 µM). Mutation of the conserved arginine residue to alanine greatly reduced the affinity to 2xS2P CTD (Kd = 8.4 ± 0.3 µM for PHF3 SPOC R1248A, 22.4 ± 2.1 µM for DIDO SPOC R1096A; Fig. 2a,b), confirming the critical role of this residue in phosphoserine recognition. SPOCD1 SPOC did not bind to unphosphorylated, single or double phosphorylated CTD peptides (Supplementary Fig. 4). This indicates that SPOCD1 SPOC may indeed have lost the phosphoserine binding ability, although we cannot exclude that it binds phosphorylated interaction partners other than the CTD.

**Fig. 2:**
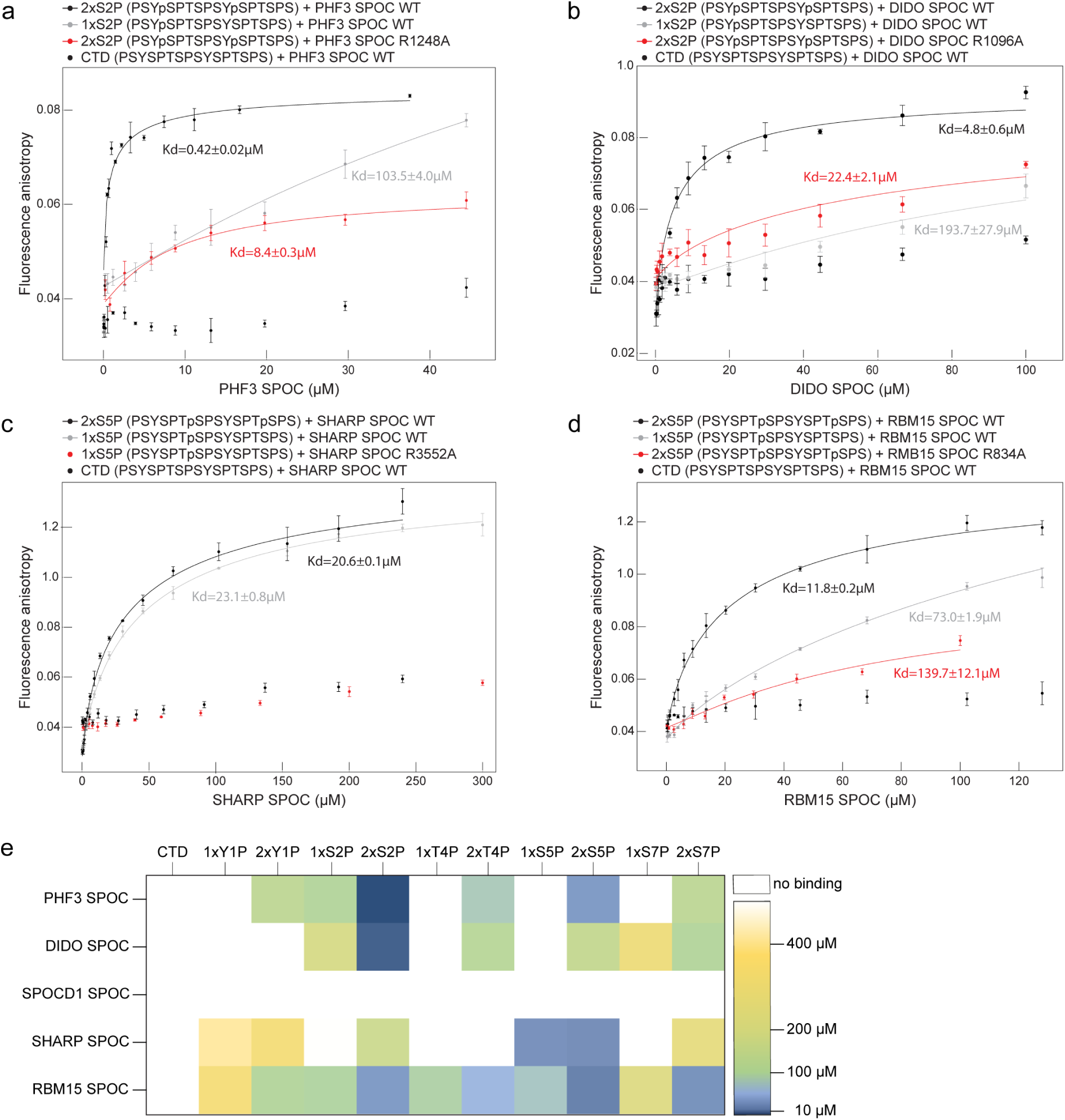
SPOC is a CTD binding domain. **a-d** Fluorescence anisotropy measurements of SPOC domains and CTD peptides. Fluorescence anisotropy is plotted as a function of protein concentration. Data points and error bars represent the mean ± standard deviation from 3 independent experiments. **e.** Heatmap of binding affinities of SPOC domains to CTD peptides.

SHARP SPOC binds to a conserved LSD-motif at the C-terminus of the transcriptional corepressors SMRT and NCoR in a serine phosphorylation-dependent manner^2, 3, 11^ (Fig. 1c, 3e). More recently, Pol II has been identified as a binding partner of SHARP SPOC^12^, supporting the idea that SHARP SPOC might be a CTD binder akin to PHF3. Indeed, SHARP SPOC bound to CTD peptides phosphorylated at serine-5 in fluorescence anisotropy assays (Fig. 2c,e). It did not bind to unphosphorylated CTD and showed very low affinity for other CTD phosphorylations (Fig. 2c,e and Supplementary Fig. 5). In contrast to PHF3 and DIDO, SHARP only required phosphorylation in a single repeat; double phosphorylation in adjacent repeats barely affected the binding affinity (Kd = 23.8 ± 0.8 µM for 1xS5P, 20.6 ± 0.1 µM for 2xS5P), which is in line with the fact that SHARP SPOC has only one conserved basic patch on its surface while PHF3 SPOC has two. Mutation of the conserved arginine residue R3552 abrogated the binding to serine-5-phosphorylated CTD (Fig. 2c).

**Fig. 3:**
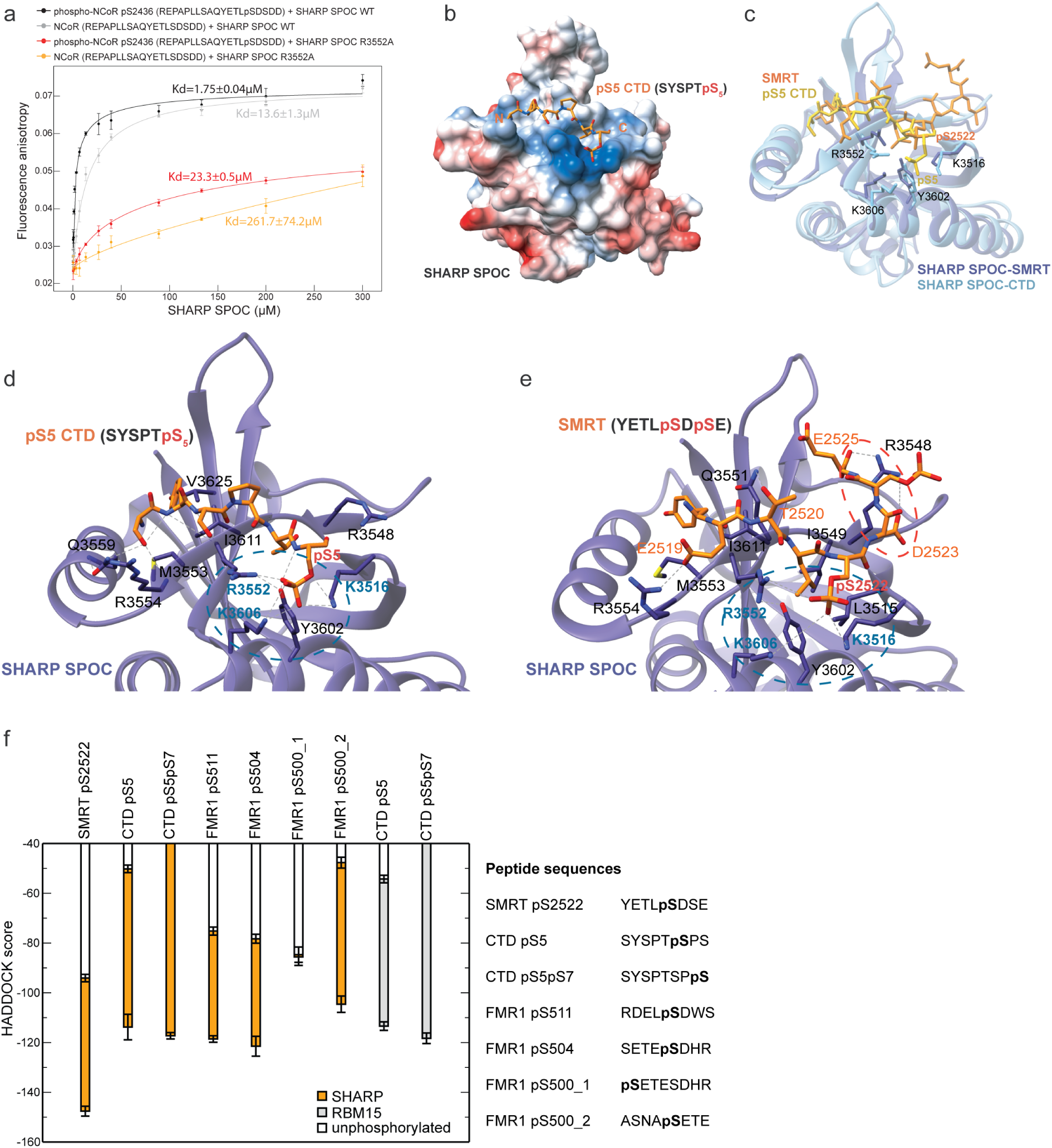
Acidic residues determine SHARP SPOC binding affinity. **a** Fluorescence anisotropy binding curves of SHARP SPOC with NCoR peptides. Fluorescence anisotropy is plotted as a function of protein concentration. Data points and error bars represent the mean ± standard deviation from 3 independent experiments. **b** Crystal structure of SHARP SPOC in complex with 1xS5P CTD peptide (7Z1K). Electrostatic surface potential was calculated using the Coulombic Surface Coloring tool in UCSF Chimera and is depicted ranging from -10 (red) to +10 (blue) kcal/(mol*e). **c** Structural comparison of SHARP SPOC binding to 1xS5P CTD and SMRT (2RT5). **d** Interactions between SHARP SPOC and 1xS5P CTD. Hydrogen bonds are indicated with dashed lines. Blue circle indicates conserved basic surface patch. **e** Interactions between SHARP SPOC and phosphorylated SMRT (2RT5). Hydrogen bonds are indicated with dashed lines. Blue circle indicates the conserved basic surface patch, red circle indicates additional electrostatic coordination of R3548 of SPOC by D2523 and E2525 of SMRT that is not possible with CTD. **f** HADDOCK scores of modeled interactions between SHARP SPOC or RBM15 SPOC and SMRT, CTD or FMR1 peptides. Peptides used for modelling were either unphosphorylated (white bars) or phosphorylated at the indicated residues (SHARP SPOC: orange bars, RBM15 SPOC: grey bars).

The SPOC domain of RBM15 interacts with the histone H3K4 methyltransferase SETD1B via an LPDSD motif similar to the SMRT/NCoR LSD motif^17^. Although it is unclear whether the serine residue in this motif is phosphorylated, K795 and K898 from the conserved basic patch are required for binding^17^, which indicates that phosphoserine binding may play a role in the interaction. In our CTD-binding analysis, RBM15 SPOC bound with highest affinity to 2xS5P CTD peptide (Kd = 11.8 ± 0.2 µM, Fig. 2d,e). Contrary to other SPOC domains, RBM15 SPOC also bound to 2xS2P and 2xS7P with similar affinities (Kd = 29.6 ± 1.4 µM for 2xS5P, 23.5 ± 0.4 µM for 2xS7P, Fig. 2e and Supplementary Fig. 6a,b), suggesting that RBM15 SPOC might be a more promiscuous phosphoserine binding domain. We modeled RBM15 SPOC binding to an 8-mer CTD peptide phosphorylated at serine-5 (Fig. 1g). In the model, phosphorylated serine-5 is tightly coordinated by the conserved residues K795, R834 and K898 (Fig. 1g). Compared to SHARP SPOC, RBM15 SPOC had a clear preference for double phosphorylated CTD peptides (Fig. 2d,e and Supplementary Fig. 6) and mutation of the highly conserved R834 to alanine reduced, but did not completely abrogate the binding to 2xS5P CTD (Fig. 2d). This indicates that there might be a second basic patch on the surface of RBM15 SPOC. Indeed, our RBM15 SPOC structure shows a potential second interaction surface at a distance from the conserved basic patch comparable to that between the two patches of PHF3 and DIDO SPOC (Fig. 1e, patch 2, distance to patch 1: 20.891 Å). Amino acids R847, R848 and K850 are potential candidates for electrostatic coordination of phosphoserine from the second CTD repeat.

In summary, SPOC domains of PHF3, DIDO, SHARP and RBM15 are CTD reader domains, which specifically recognize CTD phosphomarks via conserved basic surface patches. While PHF3, DIDO and RBM15 SPOC are tuned for binding double phosphorylated CTD-motifs, SHARP SPOC recognizes a single CTD phosphorylation.

### Acidic residues determine the binding affinity of SHARP SPOC

SHARP SPOC was reported to bind to the phosphorylated LSD motif of the SMRT/NCoR C-terminus with nanomolar affinity determined by SPR and ITC^3, 11^. To allow a direct comparison with the binding affinity to serine-5-phosphorylated CTD, we performed fluorescence anisotropy using FAM-labeled NCoR peptides that were either unphosphorylated or phosphorylated at the CK2 site S2436, which was previously shown to be critical for SHARP SPOC binding^11^ (Fig. 3a). SHARP SPOC bound to phosphorylated NCoR with an affinity of 1.75 ± 0.04 µM. Unphosphorylated NCoR had a reduced binding affinity (Kd = 13.6 ± 1.3 µM), but in contrast to previously published data the binding was not completely lost^11^. This discrepancy may be due to different experimental setups used to determine binding affinities. In line with our results, SHARP SPOC was shown to bind to the unphosphorylated C-terminus of SMRT, albeit with lower affinity than to its phosphorylated counterpart^3^.

SHARP SPOC binds to SMRT/NCoR with substantially higher affinity compared to the CTD (Kd = 1.75 ± 0.04 µM and Kd = 23.8 ± 0.8 µM, respectively) (Fig. 2c, Fig. 3a). To define the molecular determinants for the preferential binding to SMRT/NCoR, we solved the crystal structure of SHARP SPOC in complex with 1xS5P CTD peptide at a resolution of 1.55 Å (Fig. 3b,d, Supplementary Table 1). pS5 of the CTD is electrostatically anchored to the conserved basic surface patch of SHARP SPOC (Fig. 3b). The NH1 nitrogen of R3552 and the ε-aminogroups of K3516 and K3606 form hydrogen bonds with O1P, O2P and O3P of CTD pS5 (Fig. 3d). The binding mode of SHARP SPOC to 1xS5P CTD is remarkably similar to the binding between SHARP SPOC and the C-terminus of SMRT (Fig. 1c, Fig 3b-e). Both peptides occupy the same surface on the SPOC domain. pS5 of the CTD and pS2522 of SMRT are anchored via electrostatic interactions with K3516, R3552 and K3606 (Fig. 3d,e). A notable difference between the two structures are electrostatic interactions between the SMRT peptide and R3548 of SHARP SPOC, which is coordinated by D2523 and E2525 of SMRT (Fig. 3e, dashed red circle). These acidic residues are also present at the same position in the NCoR C-terminus (SMRT: - YETLpS**D**S**E**, NCoR: -YETLpS**D**S**D**D), but are absent from the CTD (-YSPTpSPSYSPTSPS) impeding hydrogen bonding with R3548 (Fig. 3d). The additional electrostatic interactions confer strong binding to SMRT even in the absence of serine phosphorylation and might explain why SHARP SPOC exhibits higher affinity for SMRT/NCoR compared to the CTD.

Given the comparable spatial proximity of SMRT residues pS2522, D2523 and E2525 and CTD residues pS5 and S7, we hypothesized that double phosphorylation of CTD at serine-5 and serine-7 might mimic the negative charge of SMRT D2523 and E2525 and confer stronger binding of SHARP SPOC to CTD. To explore this, we used HADDOCK 2.2^28, 29^ to model binding of SHARP SPOC to 8-mer SMRT pS2522, CTD pS5 and CTD pS5pS7 peptides (Fig. 3f). Based on the HADDOCK score, the strongest interaction was predicted between SHARP SPOC and phosphorylated SMRT. The predicted binding strength for double phosphorylated pS5pS7 CTD was similar to that for single phosphorylated pS5CTD, indicating that pS7 cannot compensate for the absence of acidic residues in the CTD. Similarly, RBM15 SPOC was predicted to bind CTD pS5 and CTD pS5pS7 peptides with comparable affinity (Fig. 3f).

Collectively, our findings establish SHARP SPOC as a phosphoserine binding module that preferentially binds phosphorylated LSD-motifs with adjacent acidic residues but can accommodate different phosphorylated binding partners of SHARP via its conserved basic patch.

### Transcription machinery is the major anchoring point for PHF3 and DIDO SPOC domains

To further elucidate how the SPOC domain shapes the interaction network of SPOC containing proteins, we performed co-immunoprecipitation (co-IP) of FLAG-tagged PHF3, DIDO, SHARP and RBM15 SPOC domains expressed in HEK293T cells and identified their interaction partners by mass spectrometry (Fig. 4 and Supplementary Dataset 1). The interactions were confirmed by Western blotting (Fig. 5a). Moreover, to examine SPOC-mediated interactions in the context of full-length proteins, we expressed FLAG-tagged full-length (wt) and SPOC-deleted (ΔSPOC) PHF3, DIDO, SHARP and RBM15 in HEK293T cells and performed anti-FLAG co-IP (Fig. 5b-e). All SPOC domains and full-length proteins interacted with Pol II. DIDO and PHF3 SPOC showed stronger binding compared to SHARP and RBM15, in accordance with *in vitro* binding assays (Fig. 5a and Fig. 2). The association of PHF3 with Pol II appeared weaker compared to DIDO, which may be due to the lower expression level of FLAG-tagged PHF3 SPOC (Fig. 5a). Binding of PHF3, DIDO and RBM15 to Pol II was abrogated upon loss of the SPOC domain, indicating that the SPOC-CTD interaction is of critical importance in establishing their interaction with Pol II. However, SPOC deletion did not impair interaction with Pol II, suggesting that it contacts Pol II via additional surfaces (Fig. 5e).

**Fig. 4:**
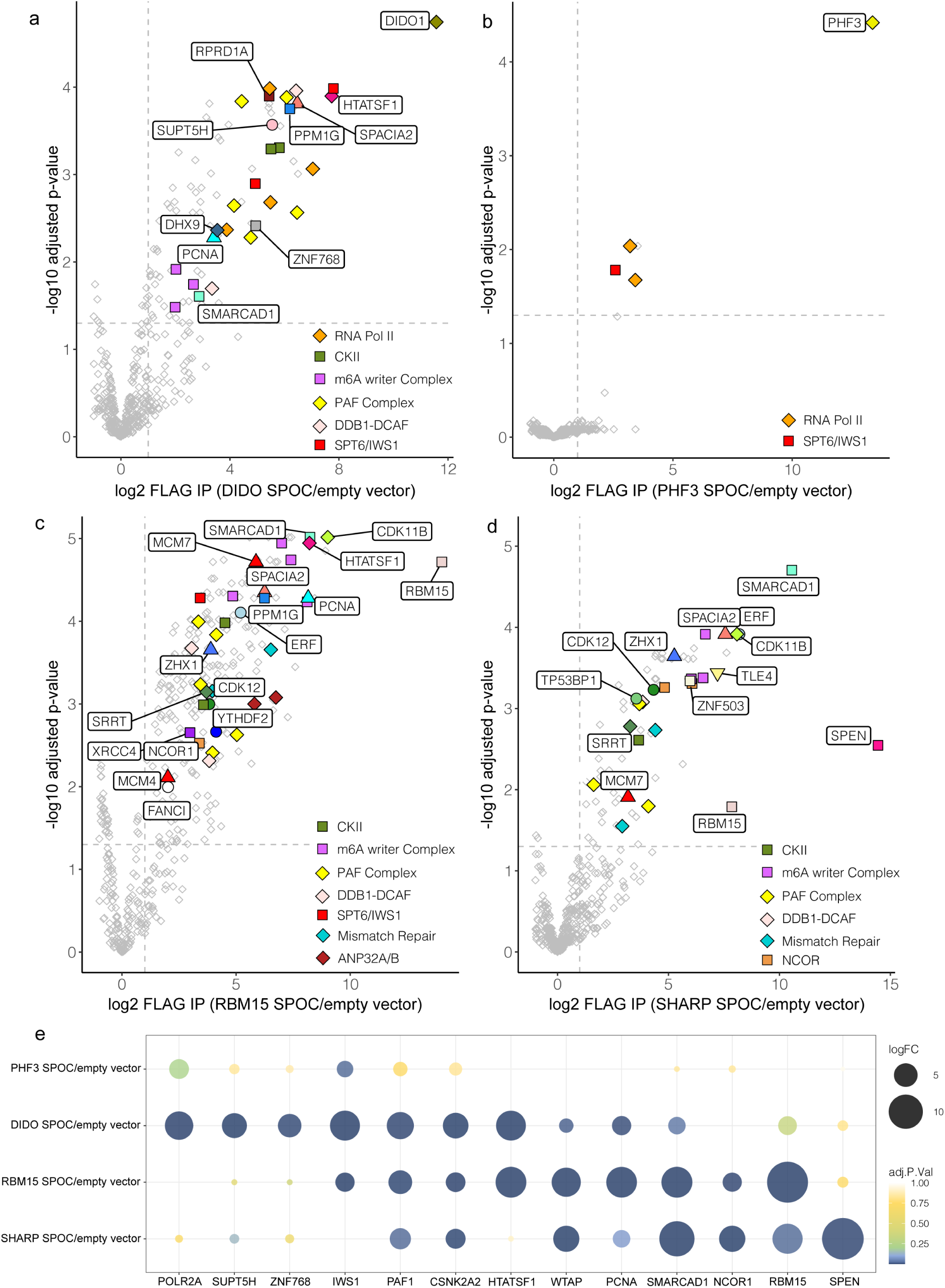
Mass spectrometry analysis of SPOC domain interactome. **a-d** Volcano plots of **a** DIDO, **b** PHF3, **c** RBM15, **d** SHARP SPOC interactors identified by mass spectrometry. **e** Overview of common SPOC domain interactors identified by mass spectrometry. The experiments were performed in three individual replicates.

**Fig. 5:**
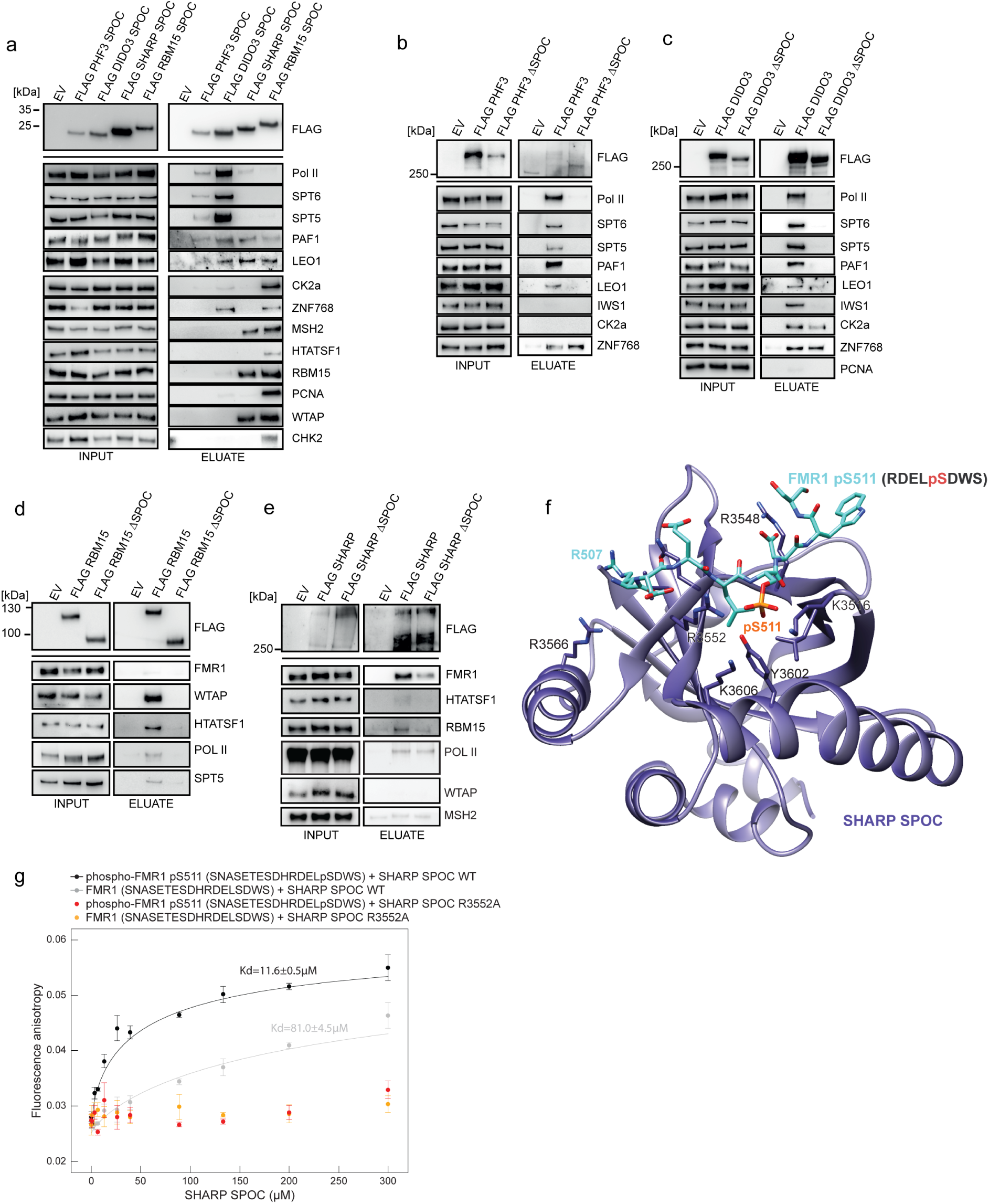
Interactome of SPOC proteins. **a** Western Blot analysis after FLAG-co-IP of FLAG-tagged SPOC domains. **b-e** Western Blot analysis after FLAG-co-IP of FLAG tagged full-length and ΔSPOC proteins. The experiments in **a-e** were performed once. **f** Structural model of the interaction between SHARP SPOC and an 8-mer FMR peptide phosphorylated at S511 generated in PyMOL and refined using the HADDOCK2.2 webserver^28, 29^. Repulsion between SHARP R3566 and FMR R507 may lead to reduced binding affinity compared to SMRT. **g** Fluorescence anisotropy binding curves of SHARP SPOC with FMR1 peptides. Fluorescence anisotropy is plotted as a function of protein concentration. Data points and error bars show mean anisotropy ± standard deviation from 3 independent experiments.

In addition to Pol II, PHF3 and DIDO showed SPOC-dependent interaction with transcription elongation factors such as SPT5, SPT6/IWS1 and the PAF1 complex (PAF1, LEO1, CTR9, CDC73, WDR61) (Fig. 4, 5a-c). Interestingly, all SPOC domains interacted with the PAF1 complex, which was not the case for full-length RBM15 and SHARP (Fig. 5a,d,e). PHF3 and DIDO interaction with the transcription factor ZNF768, which contains a heptad repeat sequence structurally related to Pol II CTD^30^ was not abrogated upon SPOC deletion (Fig. 5b,c). In contrast to PHF3, full-length DIDO and its SPOC domain showed interaction with CK2 (casein kinase 2), which was slightly diminished upon SPOC deletion (Fig. 5a,c). CK2 is known to phosphorylate a number of transcription factors, including the LSD motif of the corepressor SMRT^3, 31, 32^. The interaction between RBM15 SPOC and CK2 could not be recapitulated with the full-length RBM15 (Fig. 5a,d).

Taken together, our results establish the SPOC domain as an essential module for PHF3 and DIDO interaction with the Pol II transcription elongation machinery.

### SPOC domain mediates RBM15 and SHARP interaction with the m6A regulators

Major interactors of the RBM15 SPOC domain comprised m6A writer complex components WTAP, ZC3H13 and VIRMA/KIAA1429, the mismatch repair protein MSH2, the replication scaffold protein PCNA, as well as HTATSF1 (TAT-SF1), a transcription elongation and splicing factor that couples transcription with pre-mRNA processing^33^ (Fig. 4c and 5a). Full-length RBM15 interacted strongly with the m6A complex member WTAP and HTATSF1 in a SPOC-dependent manner but not with MSH2 or PCNA (Fig. 5a,d).

The interaction of SHARP SPOC with WTAP and MSH2 did not translate to the full-length protein (Fig. 5e). Considering the limitations of using protein overexpression and the lack of SHARP antibodies that would allow immunoprecipitation of the endogenous protein, we decided to tag SHARP and SHARP ΔSPOC with GFP at the C-terminus using CRISPR/Cas9 knock-in approach and perform anti-GFP co-IP followed by mass spectrometry (Supplementary Fig. 7a,c and 8, Supplementary Dataset 2). This analysis showed SPOC-dependent interaction of SHARP with FMR1, FXR1 and FXR2, which was confirmed for FMR1 by Western blotting (Fig. 5e and Supplementary Dataset 2). FMR1, also called FMRP (fragile X mental retardation protein), binds to m6A mRNA and mediates its export into the cytoplasm, which is essential for neuronal differentiation^34^. Both FXR1 and FXR2 (fragile X-related proteins 1 and 2) regulate adult neurogenesis and have been implicated in various neurological disorders, but the underlying mechanisms are not well understood^35^. Given that SHARP SPOC binds to LSD motifs at the C-terminus of SMRT/NCoR^3, 11^, we hypothesized that SHARP may contact FMR1 via a similar motif. We identified an LSD motif with several phosphorylated residues based on PhosphoSitePlus^36^ and used HADDOCK 2.2^28, 29^ to refine SHARP/FMR1 complexes, which were reconstructed using the SHARP-SMRT structure (2RT5) as a template (Fig. 3f). According to the modeling, SHARP interactions with FMR1 are weaker compared to SMRT. This can be due to the absence of a tyrosine residue at the corresponding position (Y2518 in SMRT or Y1 in CTD) that interacts with SHARP R3566 via π-π interactions and forms hydrophobic contacts with M3553. In the FMR pS511 peptide the residue in this position is arginine (R507) instead of tyrosine that can lead to repulsive interaction with R3566 of SHARP SPOC (Fig. 5f).

We determined the binding affinity of SHARP SPOC to the peptide surrounding pS511 by fluorescence anisotropy (Fig. 5g). SHARP SPOC bound to the FMR1 pS511 peptide with an affinity of 11.6 ± 0.5 µM, further showing how the SHARP SPOC domain mediates interaction with a variety of phosphoserine binding partners. Affinity for the unphosphorylated peptide was strongly reduced (Kd = 81.0 ± 4.5 µM), while mutation of the conserved R3552 to alanine abrogated the binding to both peptides.

Overall, our results suggest that the SPOC domains of the Spen family members RBM15 and SHARP mediate interactions with the m6A writer and reader proteins.

### SPOC determines genomic localization of PHF3 and SHARP

SPOC proteins regulate transcription and co-transcriptional processes. Thus, their recruitment to chromatin is crucial for their proper function. To evaluate the importance of the SPOC domain for their genomic localization we employed Chromatin Immunoprecipitation followed by high-throughput sequencing (ChIP-seq) analysis of SHARP and PHF3 (Fig. 6). Due to a lack of suitable antibodies for immunoprecipitation of the endogenous proteins, we performed GFP-ChIP on endogenously tagged SHARP-GFP, SHARP ΔSPOC-GFP, PHF3-GFP and PHF3 ΔSPOC-GFP cell lines. We had previously generated PHF3-GFP and PHF3 ΔSPOC cell lines^5^ and applied CRISPR/Cas9 knock-in to tag PHF3 ΔSPOC with GFP at the C-terminus (Supplementary Fig. 7b,d). ChIP-seq analysis revealed that PHF3-GFP bound to expressed genes and that recruitment to chromatin was reduced upon deletion of the SPOC domain. Among PHF3 target genes were *BEX5* and *HOXA5* (Fig. 6a,b), We confirmed binding of PHF3-GFP to these genomic loci by ChIP-qPCR (Fig. 6a,b). Both in sequencing and qPCR analysis, PHF3 recruitment was strongly reduced upon deletion of the SPOC domain.

**Fig. 6:**
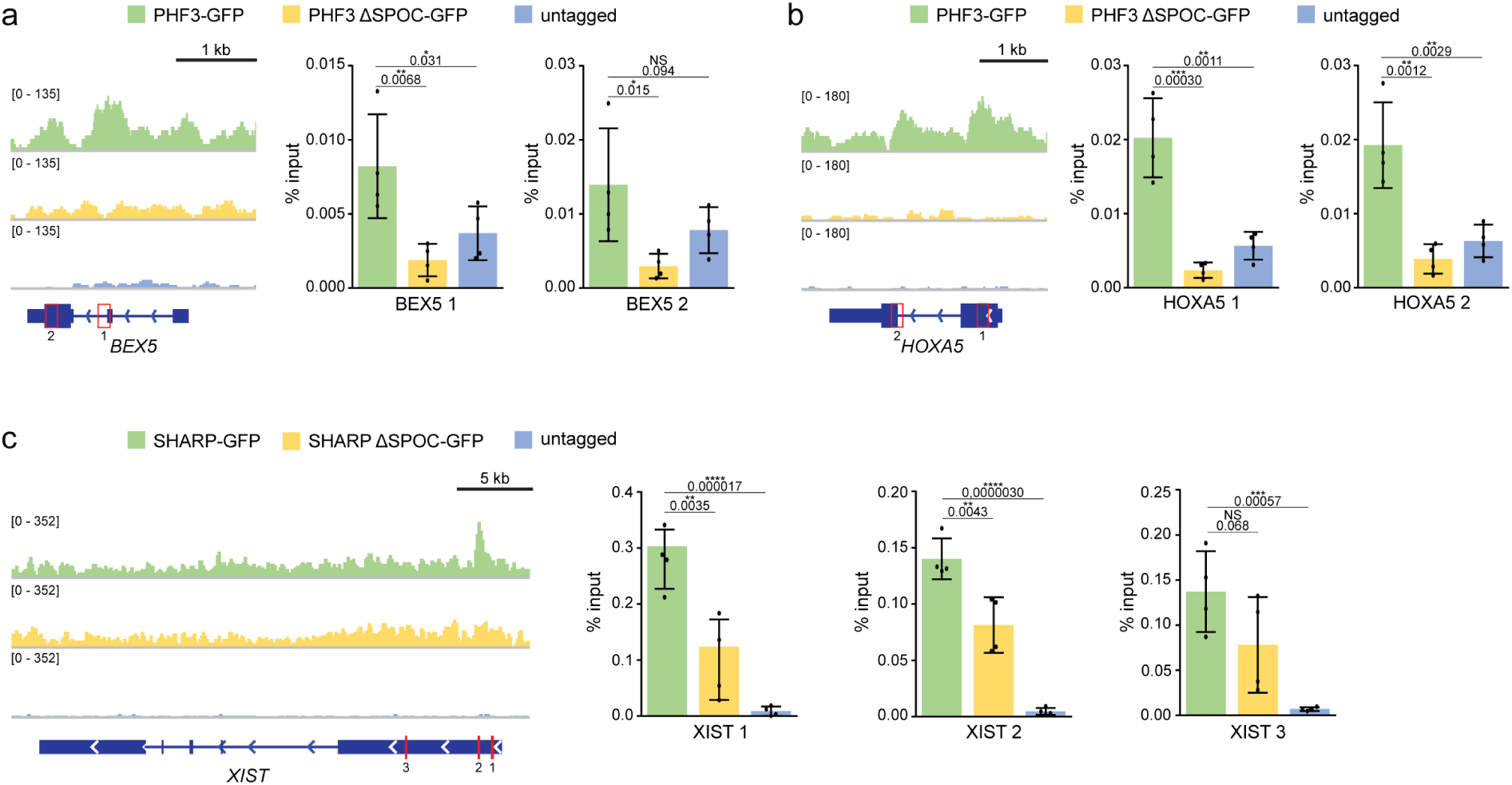
SPOC domains determine genomic localizaton. **a** Integrative Genomics Viewer (IGV) snapshots showing GFP-ChIP-seq reads and GFP-ChIP-qPCR analysis for BEX5 in PHF3-GFP, PHF3 ΔSPOC-GFP and untagged HEK 293T cells. **b** IGV snapshots showing GFP-ChIP-seq reads and GFP-ChIP-qPCR analysis for HOXA5 in PHF3-GFP, PHF3 ΔSPOC-GFP and untagged HEK 293T cells. **c** IGV snapshots showing GFP-ChIP-seq reads and GFP-ChIP-qPCR analysis for XIST in SHARP-GFP, SHARP ΔSPOC-GFP and untagged HEK 293T cells. The qPCR data in **a-c** are presented as mean ± standard deviation of four individual experiments, individual data points are indicated as black dots. One-tailed, two-sample equal variance t-test was used to determine p-values. qPCR amplicons are indicated as red boxes.

ChIP-seq analysis of SHARP-GFP revealed only one area of enriched genomic binding at the *XIST*-locus in HEK293 cells, which we confirmed by qPCR using multiple primer pairs targeting the *XIST* locus (Fig. 6c). Similar to PHF3, SHARP ΔSPOC was less strongly recruited to chromatin than the full length protein (Fig. 6c).

Our results indicate that SPOC proteins are recruited to chromatin via the SPOC domain, further highlighting the important role of this domain in SPOC protein-mediated gene regulation.

## Discussion

We established SPOC as a CTD reader domain that recognizes different CTD phosphorylation patterns. PHF3 and DIDO SPOC bind to CTD phosphorylated at serine-2, while SHARP SPOC binds phosphorylated serine-5. RBM15 SPOC has a preference for serine-5 phosphorylation, but also binds phosphorylated serine-2 and serine-7 with similar affinity. serine-5 and serine-7 phosphorylation are prominent in early stages of transcription and decrease during the elongation phase concurrent with an increase of serine-2 phosphomarks^18^. This indicates that SHARP binds to Pol II during early transcription, PHF3 and DIDO regulate the elongation stage, while RBM15 might function in different phases of transcription (Fig. 7).

**Fig. 7:**
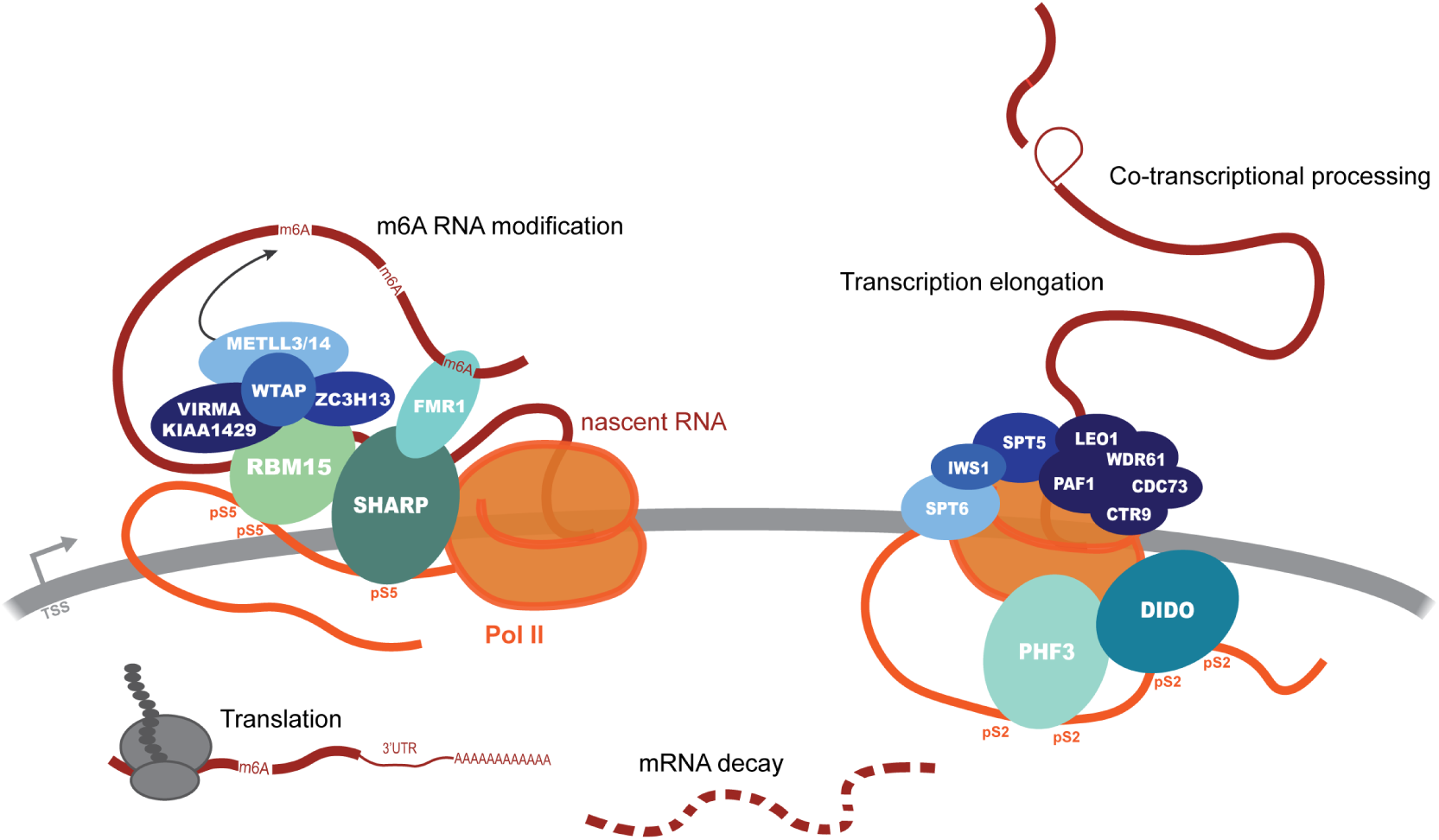
SPOC bridges transcription with co-and post-transcriptional processes: SHARP and RBM15 interact with RNA and writers and readers of m6A RNA modification, while PHF3 and DIDO interact with regulators of transcription and co-transcriptional processing. SPOC protein interactors in turn regulate downstream processes like translation and mRNA decay.

DIDO SPOC, like the SPOC domain of its paralogue PHF3, preferentially binds to CTD phosphorylated on serine-2 in two adjacent repeats. The SPOC domain is essential for interaction with Pol II. Interestingly, the SPOC domain is only present in the longest isoform DIDO3; the short isoform DIDO1 does not contain SPOC while DIDO2 SPOC domain is truncated and likely unfunctional. Since association with Pol II and the elongation complex is dependent on the SPOC domain, direct interaction of DIDO1 and DIDO2 with Pol II is improbable. DIDO3 is the dominant isoform in embryonic stem cells and promotes maintenance of the stem cell state; a switch to expression of the short isoform DIDO1 triggers differentiation^22^. While DIDO3 is expected to regulate transcription through direct interaction with Pol II, the shorter isoforms may indirectly modulate gene expression through chromatin binding.

*In vitro* binding assays showed that DIDO SPOC binds to 2xS2P CTD with 10-fold lower affinity compared to PHF3 SPOC (Fig. 2a,b). However, co-IP experiments with SPOC domains or full-length proteins show that DIDO may have similar or even stronger affinity for Pol II in cells, which may be due to higher expression levels and stability compared to PHF3 (Fig. 5a-c). While DIDO and PHF3 dock onto the Pol II elongation complex through interaction with CTD phosphorylated at serine-2, they can establish additional contacts with Pol II, for instance through their TLD domain which has been shown to bind to the Pol II jaw domain in their yeast homologue Bye1^37^. The TLD shares homology with the central domain of transcription factor IIS but lacks the ability to rescue backtracked Pol II by stimulating RNA cleavage, which is conferred by TFIIS domain III^38^. PHF3 can outcompete TFIIS for Pol II binding and thereby repress transcription^5^. It remains to be addressed in future studies if DIDO regulates transcription in a similar fashion and whether PHF3 and DIDO share regulatory functions. Given that both PHF3 and DIDO bind to the same CTD phosphomark, it is conceivable that they might either bind to the elongation complex simultaneously and exercise synergistic functions or that their binding might be mutually exclusive and they compete for Pol II binding. Given their important roles in neuronal development^5, 9^ they might also act redundantly. While genomic localization of PHF3 is largely dependent on the SPOC domain, DIDO PHD, in contrast to PHF3 PHD, binds to H3K4me3 histone marks and can thus bind chromatin independently of SPOC^39, 40^. DIDO may therefore have independent functions that cannot be taken over by PHF3.

It was previously shown that SHARP SPOC interacts with both SMRT/NCoR containing corepressor complexes and KMT2D containing activator complexes, with phosphorylation of the SMRT/NCoR LSD motif shifting the balance to the repressive complex^11^. Our study adds new members to the SHARP SPOC interactome. We showed that SHARP SPOC binds to Pol II CTD phosphorylated on serine-5, albeit with lower affinity than to SMRT/NCoR. We solved the structure of SHARP SPOC in complex with the CTD, which revealed that the strength of binding depends in part on an acidic residue adjacent to the LSD motif of SMRT/NCoR, which is missing in the CTD. Although co-IP experiments showed that the SPOC domain is not essential for the interaction between SHARP and Pol II (Fig. 5e), ChIP analysis revealed reduced recruitment of SHARP ΔSPOC onto the Xist locus (Fig. 6). This suggests a bimodal recruitment of SHARP onto the Xist locus on the X-chromosome through direct interactions between SHARP RRMs and Xist lncRNA^41–43^ and between SHARP SPOC and Pol II CTD (Fig. 2c). SHARP RRMs and SPOC domain are essential for Xist lncRNA-mediated silencing during X-chromosome inactivation^12^. Although initially suggested to act via recruiting HDAC3, Xist-mediated silencing is abolished by SHARP deletion, but only attenuated upon HDAC3 deletion^44, 45^, suggesting that additional mechanisms are involved in SHARP-dependent repression of the X-chromosome. One of the earliest events upon Xist upregulation at the onset of X-chromosome inactivation is exclusion of Pol II from the chromosome territory^46^, but the underlying mechanism of Pol II exclusion remains unknown. Our finding that SHARP directly interacts with Pol II CTD opens up the intriguing possibility that SHARP might initiate gene silencing by directly acting on Pol II.

We furthermore identified FMR1 (FMRP) as a SPOC-dependent interactor of SHARP. An expansion of CGG-repeats within FMR1 leads to reduction or abolishment of its expression and is the cause of fragile X syndrome, which can manifest as mild to severe intellectual disability and autism spectrum disorder^47^. On a molecular level, FMR1 is an RNA-binding protein that binds m6A mRNA, regulates mRNA export and acts as a translational repressor to modulate the amount of protein production^34, 48^. We show that SHARP SPOC binds to an LSD motif in FMR1 in a phosphorylation-dependent manner. The physiological role of this interaction is yet to be resolved; however, SHARP and its SPOC domain have also been implicated in neurodevelopment^7, 8^ and SHARP and FMR1 might cooperate to ensure proper formation of neurons with SHARP controlling the transcriptional aspects and FMR1 regulating mRNA export and translation.

Interaction with the m6A regulators may be a common theme for the Spen family of SPOC proteins. While SHARP interacts with the m6A reader protein FMR1 in a SPOC-dependent manner, RBM15 SPOC is crucial for its interaction with the m6A writer complex (Fig. 5d,e). RBM15 was previously shown to recruit the m6A writer complex to RNA via WTAP and ZC3H13, but the mechanism remained unclear^16^. Our results suggest that RBM15 uses its SPOC domain to recruit the m6A machinery onto RNA bound by RBM15 RRMs (Fig. 7). The m6A RNA modification can regulate mRNA decay and translation^49^, raising the question as to the importance of RBM15 SPOC for the regulation of mRNA levels.

Analogous to PHF3- and DIDO-mediated co-regulation of transcription through SPOC- dependent binding to Pol II CTD, RBM15 and SHARP may cooperate in co- and post-transcriptional RNA processing through m6A RNA modification. Interestingly, SHARP interacts with RBM15 in a SPOC-dependent manner (Fig. 5a,e), suggesting that SHARP and RBM15 may work together to recruit the m6A writer complex onto target RNAs such as Xist lncRNA (Fig. 7). Xist RNA is modified with m6A, which contributes to Xist-mediated X-chromosome silencing^50^. Knock-out of RBM15 or m6A writer complex components shows reduced, but not completely abrogated Xist m6A and a modest effect on X-chromosome silencing^45^. The role of SHARP in the regulation of Xist m6A modification remains to be further explored.

Taken together, we showed that SPOC domains from PHF3, DIDO, SHARP and RBM15 have a different affinity and specificity towards CTD phosphomarks encoded in their distinct surface electrostatic potential patterns, suggesting that SPOC is a versatile Pol II CTD reader. Moreover, SPOC domains are versatile phosphoserine readers engaging with the Pol II transcription elongation complex, co-repressor complexes, m6A writer and reader machineries. Multivalent interactions of SPOC domain proteins facilitate coupling between transcription and RNA metabolism to ensure appropriate gene expression.

## Materials and Methods

### Cloning

For mammalian expression, human DIDO3, SHARP and RBM15 were amplified from HEK293T cDNA and cloned into CMV10 N3XFLAG (Sigma) by Gibson assembly (NEB). These constructs were used for cloning ΔSPOC truncations using Gibson assembly (NEB). CMV10 N3XFLAG-PHF3 and PHF3 ΔSPOC constructs were generated previously^5^. NLS-SPOC constructs were cloned into CMV10 N3XFLAG (Sigma) between NotI and XbaI. NLS sequence (RAPKKKRKVGG) was introduced to ensure nuclear localization of the isolated SPOC domains. For bacterial expression, human DIDO3 SPOC (1047-1205aa), SHARP SPOC (3496-3664aa), RBM15 SPOC (775-960aa) and SPOCD1 SPOC (858-1025aa) were cloned into pET M11 between NcoI and XhoI for N-terminal His6 fusion. His6-PHF3 SPOC was generated previously^5^. Arginine mutations were introduced by site-directed mutagenesis according to the FastCloning protocol^51^. Repair templates for CRISPR/Cas9 knock-in were cloned by Gibson assembly. Primer sequences are listed in Supplementary Table 3.

### Protein purification

SPOC domains were expressed in *E.coli* Rosetta2 (DE3) cells (Novagen). O/N cultures were diluted 1:50 in 2xTY, grown at 37°C until OD_600_=0.8 and induced with 0.5 mM IPTG for 3h at 30°C. His_6_- tagged SPOC domains were purified by affinity chromatography using HisTrap HP column (Cytiva) or His-Pur Ni-NTA resin (Thermo Scientific) equilibrated in 25 mM Tris-Cl pH 7.4, 500 mM NaCl, 20 mM Imidazole. Elution with 25 mM Tris-Cl pH 7.4, 500 mM NaCl, 500 mM Imidazole was followed by TEV cleavage of the His_6_-tag and size exclusion chromatography using Sephacryl S-200 HR 16/600 or 26/600 (Cytiva) equilibrated in 25 mM Tris-Cl pH 7.4, 25 mM NaCl, 1 mM DTT.

### Peptide labeling

CTD peptides were N-terminally labelled with Atto-488 while NCoR and FMR1 peptides were purchased with an N-terminal FAM-label. 0.5 mg of lyophilized CTD peptides were dissolved in 30 µl DMSO (Sigma-Aldrich). One molar equivalent of Atto-488 NHS ester (Atto-tec) and five equivalents DIPEA (Sigma-Aldrich) were added to three equivalents of peptide. Reactions were incubated ON at room temperature protected from light. Labeled peptides were purified by reverse-phase HPLC over a C18 column (Agilent Technologies) using a gradient from 30 to 70 % Methanol (Merck). Purified peptides were lyophilized and stored at -20°C.

### Fluorescence anisotropy

Fluorescence anisotropy measurements were performed on a Perkin Elmer LS50 B fluorescence spectrometer operated by Perkin Elmar FL WinLab software (version 3.00). Excitation wavelength was set to 500 nm (excitation slit 10 nm), emission wavelength to 518 nm (emission slit 6 nm), integration time to 1 sec. Measurements were conducted at room temperature in 25 mM Tris-Cl pH 7.4, 25 mM NaCl, 1 mM DTT. The grating factor was determined using unconjugated Atto-488 dye. 90 nM peptide were titrated with increasing concentrations of SPOC domains in a final volume of 110 µL. Each individual datapoint is an average of 20 measurements. For each binding curve, datapoints were acquired from three independent SPOC dilution series. Data are presented as mean anisotropy ± standard deviation from the three independent replicates. Data were plotted and fitted with QtiPlot 1.0.0-rc13 software (version 5.9.8.)

### X-ray crystallography

Crystalization was performed at at 22°C or 4°C using a sitting-drop vapour diffusion technique and micro-dispensing liquid handling robot Mosquito (TTP labtech). The best diffracting crystals of SHARP SPOC:1xS5P CTD were grown at 22°C in conditions E9 from ShotGun HT screen (SG1 HT96 Molecular Dimensions, Suffolk, UK) containing 0.1 M Potassium thiocyanate, 30% w/v PEG 2000 MME at protein concentration 10 mg/ml. The best diffracting crystals of RBM15 SPOC were grown at 4°C in conditions F12 from PACT screen (PACT premier HT96 Molecular Dimensions, Suffolk, UK) containing 0.2 M Sodium malonate dibasic monohydrate, 0.1 M Bis-Tris propane pH 6.5, 20% w/v PEG 3350 at protein concentration 6 mg/mL. The crystals were flash cooled in liquid nitrogen prior to data collection. The data set of SHARP SPOC:1xS5P CTD was collected at the beamline MASSIF beamline ID30a1 (ESRF, Grenoble) at 100K using a wavelength of 0.966 Å. The data set of RBM15 was collected at the I04 (DLS, Didcot Oxfordshire) 100K using a wavelength of 0.9795 Å. The data frames were processed using the XDS package^52^, and converted to mtz format with the program AIMLESS^53^ with the help of autoprocessing pipelines at the synchrotrons. The structures were determined using the molecular replacement program PHASER with atomic coordinates of SPOC SHARP (PDB code 1OW1) as a search model in case of SHARP SPOC:1xS5P structure and our SHARP SPOC structure as a search model for RBM15. The structures were then refined with Phenix Refine v 1.20.1-4487^54^ and rebuilt using Coot v 0.9.6^55^.

### Modeling of SPOC complexes with peptides

3D configurations of complexes of SHARP SPOC with all peptides in phosphorylated and unphosphorylated forms (except the CTD fragment) were generated using PyMOL starting from the SHARP-SMRT structure (2RT5)^3^ as a template. The X-ray structures refined in the present study were used as templates for complexes involving phosphorylated CTD 8-mer repeat (SYSPTSPS) and SHARP and RBM15 SPOC domains, whereby the CTD peptide was extended to 8 residues. The structure of RBM15 SPOC predicted by AlphaFold (see https://alphafold.ebi.ac.uk/entry/Q96T37) was also used for this purpose. All complexes were further refined using HADDOCK 2.2 web-server ^28, 29^. The peptide length was fixed to 8 residues in all cases. The HADDOCK score, which was calculated after model refinement for different complexes, was used as a proxy for stability (Fig. 3f).

### Cell Culture

HEK293T cells were grown in Dulbecco’s Modified Eagle’s Medium (DMEM 4.5 g/L glucose) (Sigma) supplemented with 10% fetal bovine serum (Sigma), 1% L-glutamine (Sigma), 1% penicillin-streptomycin (Sigma) under 5% CO2 at 37 °C. To generate endogenously tagged SHARP-GFP and SHARP ΔSPOC-GFP cell lines by CRISPR/Cas9, gRNAs targeting the 3’end of *SHARP* (SHARP 3’gRNA: 5’-CCACTCAGTGGCTCACACGG-3’) and the region upstream of SPOC (SHARP SPOC gRNA: 5’-GAACCATATCCACGGGTCTC) were designed and cloned into pX330 plasmid encoding Cas9 nuclease^56^. HEK 293T cells were transfected with 2 µg pX330-SHARP 3’gRNA and 4 µg plasmid repair template comprising EGFP-P2A-puromycin and flanking ≈1kb homology arms for SHARP-GFP or 2 µg pX330-SHARP 3’gRNA + 2 µg pX330-SHARP SPOC gRNA to excise the region encoding SPOC and 4 µg plasmid repair template for SHARP ΔSPOC-GFP. 2 days after transfection 0.5 µg/µL puromycin was added to the culture medium for 1 week. To allow for recovery surviving cells were grown for several days without puromycin. GFP-positive cells were sorted into 96-well plates by FACS. Colonies originating from single cells were expanded, genomic DNA was extracted, positive clones were identified by PCR and sequencing. To generate endogenously tagged PHF3 ΔSPOC-GFP cell line, HEK 293T PHF3 ΔSPOC cells were transfected with 2 µg each of pX335 plasmids encoding Cas9 nickase and gRNAs targeting the *PHF3* 3’end (5’-CAGTGTGGTCCCTATCTTTG-3′ and 5′-TAAAATTTGCAGGCTGCTTC-3) and 4µg plasmid repair template containing EGFP-P2A-puromycin and flanking 1.5 kb homology arms. The PHF3 ΔSPOC cell line and plasmids had been generated for a previous study^5^. Selection and identification of positive clones was performed as described above.

### Immunoprecipitation

For anti-FLAG immunoprecipitation, HEK293T cells were transfected with the following FLAG constructs: 3xFLAG-NLS-PHF3 SPOC, 3xFLAG-NLS-DIDO3 SPOC, 3xFLAG-NLS-SHARP SPOC, 3xFLAG-NLS-RBM15-SPOC, 3xFLAG-PHF3, 3xFLAG-PHF3 ΔSPOC, 3xFLAG-SHARP, 3xFLAG-SHARP ΔSPOC, 3xFLAG-DIDO3, 3xFLAG-DIDO3 ΔSPOC, 3xFLAG-RBM15, 3xFLAG-RBM15 ΔSPOC. Cells were seeded on a 10 cm dish one day prior to transfection, transfected with 8 µg of the respective plasmid at 60-70% confluency and harvested 48 hours after transfection. For anti-GFP immunoprecipitation, one 10 cm dish of HEK293T cell lines expressing GFP-tagged SHARP WT or ΔSPOC was used per immunoprecipitation. Pellets were lysed in 1 ml lysis buffer (50 mM Tris-Cl pH 8, 150 mM NaCl, 0.1% Triton, 1x protease inhibitors, 2mM Na_3_VO_4_, 1 mM PMSF, 2mM NaF, 50 units/ml benzonase and 1 mM DTT) for 30 min on a rotating wheel at 4°C. Protein concentrations were measured using the Bradford protein assay using NanoDrop 2000c. The volume used for IP was adapted to the lysate with the lowest measured protein concentration to ensure that the same amount of protein lysate was used for immunoprecipitation. Lysates were incubated for 2 h on a rotating wheel at 4°C with anti-FLAG M2 beads (Sigma). Beads were washed once with lysis buffer (without benzonase) and 4 times in TBS. For mass spectrometry analysis the samples were further processed as described in Mass spectrometry sample preparation. For western blot analysis of anti-FLAG immunoprecipitation, the beads were eluted with 30 µl of 3xFLAG peptide (150 ng/µl) in TBS for 30 min on a rotating wheel at 4°C. 10µl of the 4xSDS loading buffer was added to the eluate and half of the total volume (20 µl) was analysed by SDS-PAGE followed by western blotting. Antibodies are listed in Supplementary Table 4.

### Mass spectrometry sample preparation

Magnetic anti-FLAG beads were transferred to fresh tubes and resuspended in 50 µl of 50 mM ammonium bicarbonate (ABC). The proteins were digested with 200 ng Lys-C (WAKO) at 37 °C shaking at 1200 rpm for 3 hours. The supernatant was transferred to a fresh tube (L-fraction). Still bound proteins were eluted from the beads by adding 20 µL of 100 mM glycine pH 2. After gentle vortexing and incubation at RT for 2-5 min, the supernatant was transferred to a fresh tube (G-fraction). Elution step was performed three times in total, the eluates were combined and the pH was made alkaline using 1 M Tris-HCl pH 8. L- and G- fractions were subsequently processed in parallel. Disulfide bonds were reduced with 10 mM dithiothreitol for 45 min at 56 °C. Alklyation was performed with 20mM iodoacetamide at room temperature for 45 min in the dark. Remaining iodoacetamide was quenched by adding 5 mM DTT and the proteins were digested with 200 ng trypsin (Trypsin Gold, Promega) at 37 °C ON. The digest was stopped by addition of trifluoroacetic acid (TFA) to a final concentration of 0.5 %, and the peptides were desalted using C18 Stagetips. After quality check of the digests by liquid chromatography, L- and G-fractions from the respective sample were combined. Beads with cross-linked anti-GFP nanobody were transferred to fresh tubes and resuspended in 30 µL of 2 M urea in 50 mM ammonium bicarbonate (ABC). Disulfide bonds were reduced with 10 mM dithiothreitol for 30 min at room temperature before adding 25 mM iodoacetamide and incubating for another 30 min at room temperature in the dark. The remaining iodoacetamide was quenched by adding 5 mM DTT and the proteins were digested with 150 ng trypsin (Trypsin Gold, Promega) for 90 min at room temperature in the dark. The supernatant was transferred to a fresh tube. The beads were rinsed with 30 µL of 2 M urea in 50 mM ABC. The combined supernatants were diluted to 1 M urea with 50mM ABC, additional 150 ng trypsin (Trypsin Gold, Promega) were added and incubated at 37 °C ON. The digest was stopped by the addition of 10 % trifluoroacetic acid (TFA) to a final concentration of 0.5 %, and the peptides were desalted using C18 Stagetips^57^.

### Liquid chromatography-mass spectrometry

Peptides were separated on an Ultimate 3000 RSLC nano-HPLC system using a pre-column for sample loading (Acclaim PepMap C18, 2 cm × 0.1 mm, 5 μm), and a C18 analytical column (Acclaim PepMap C18, 50 cm × 0.75 mm, 2 μm, all HPLC parts Thermo Fisher Scientific), applying a linear gradient from 2% to 35% solvent B (80% acetonitrile, 0.1 % formic acid; solvent A 0.1 % formic acid) at a flow rate of 230 nL/min over 120 min. The GFP-IPs were analyzed on a Q Exactive HF-X Orbitrap and the FLAG-IPs on an Orbitrap Fusion Lumos mass spectrometer coupled to the HPLC via the EASY-Spray ion-source (all Thermo Fisher Scientific) equipped with coated emitter tips (New Objective). The mass spectrometers were operated in data-dependent acquisition mode (DDA). On the Q Exactive HF-X survey scans were obtained in a mass range of 375-1500 m/z with lock mass off, at a resolution of 120000 at 200 m/z and an AGC target value of 3E6. The 8 most intense ions were selected with an isolation width of 1.6 m/z, for max. 250 ms at a target value of 1E5, and then fragmented in the HCD cell at 28 % normalized collision energy. Spectra were recorded at a resolution of 30000. Peptides with a charge of +1 or >+6 were excluded from fragmentation, the peptide match feature was set to preferred, the exclude isotope feature was enabled, and selected precursors were dynamically excluded from repeated sampling for 30 seconds. On the Orbitrap Lumos Fusion the survey scans were obtained in a mass range of 375-1500 m/z, at a resolution of 120000 at 200 m/z, with an AGC target value of 4E5. In a cycle time window of 2.5 s, the most intense precursors were selected with an isolation width of 1.0 m/z, aiming at an AGC target value of 2E5 within 150 ms, and fragmented in the HCD cell with a collision energy of 30 %. The spectra were recorded in the orbitrap at a resolution of 30000. Peptides with a charge of +2 to +6 were included for fragmentation, the MIPS mode was set to “peptide” and the exclude isotope feature was enabled. Selected precursors were dynamically excluded from repeated sampling for 45 seconds.

### Mass spectrometry data analysis

Raw data were processed using the MaxQuant software package^58^ (version 1.6.16.0 respect. 1.6.14.0) searching against the Uniprot human reference proteome (January 2020, www.uniprot.org) as well as a database of most common contaminants. The search was performed with full trypsin specificity and a maximum of two missed cleavages. Carbamidomethylation of cysteine residues was set as fixed, oxidation of methionine, and acetylation of protein N-termini as variable modifications. For the FLAG-IP samples, acetylation of lysines and phosphorylation of serine, threonine and tyrosine were additionally defined as variable modifications. For label-free quantification the “match between runs” feature and the LFQ function were activated - all other parameters were left at default. Results were filtered at a false discovery rate of 1% at protein and peptide spectrum match level. MaxQuant output tables were further processed in R^59^ (version 4.0.2). Reverse database identifications, contaminant proteins, protein groups identified only by a modified peptide, protein groups with less than two quantitative values in one experimental group, and protein groups with less than 2 razor peptides were removed for further analysis. Missing values were replaced by randomly drawing data points from a normal distribution modeled on the whole dataset (data mean shifted by -1.8 standard deviations, width of the distribution of 0.3 standard deviations). Differences between groups were statistically evaluated using the LIMMA package^60^ at 5% FDR (Benjamini-Hochberg).

### Chromatin Immunoprecipitation

Cells were resuspended in 50 mL PBS/10^8^ cells and fixed by adding 1% formaldehyde for 10 min. Formaldehyde was quenched by addition of 0.6 M glycine pH 3 for 15 min. Cells were washed twice in cold PBS. Nuclei were isolated by resuspending the cells in 5 mL cold lysis buffer 1 (50 mM HEPES/KOH pH 7.5, 140 mM NaCl, 1 mM EDTA, 10% glycerol, 0.5% Nonidet P-40, 0.25% Triton X-100, 1x Complete protease inhibitors (Roche)) per 10^8^ cells and rotating at 4°C for 10 min. After centrifugation nuclei were resuspended in 5mL/10^8^ cells cold lysis buffer 2 (10 mM Tris-Cl pH 8, 200 mM NaCl, 1 mM EDTA, 0.5 mM EGTA, 1x Complete protease inhibitors (Roche)) and rotated for 10 min at room temperature. The pellet after centrifugation was resuspended in 3 mL lysis buffer 3 (10 mM Tris-Cl pH 8, 100 mM NaCl, 1 mM EDTA, 0.5 mM EGTA, 0.1% Na-deoxycholate, 0.5% N-lauroylsarcosine, 1x Complete protease inhibitors (Roche)) and chromatin was sheared to an average size of 200-600 bp by sonication for 20 cycles 30 sec on/30 sec off using Bioruptor Pico (Diagenode). 1% Triton X-100 was added after sonication. 5-10% of chromatin was kept as an input control for qPCR. For ChIP-seq 1.5% spike-in chromatin from a mouse cell line expressing endogenously tagged PHF3-GFP was added. Anti-GFP antiserum (Abcam ab290) was added to sheared chromatin and rotated ON at 4°C. Protein A Dynabeads (Invitrogen) were washed in cold block solution (0.5% BSA in PBS) three times, mixed with antibody-bound chromatin and rotated for 4-6 h at 4°C. Beads were washed in RIPA washing buffer (50 mM Hepes/KOH pH 7.5, 500 mM LiCl, 1 mM EDTA, 1% NP-40, 0.7% Na-deoxycholate) five times and in 50 mM NaCl in TE once. Immunoprecipitated protein-DNA complexes were eluted in 200 µL elution buffer (50 mM Tris-Cl pH 8, 10 mM EDTA, 1% SDS) for 15 min at 65°C. Eluates were incubated ON at 65°C to reverse crosslinks, treated with 0.2 mg/mL RNase A for 2 h at 37°C and 0.2 mg/mL proteinase K and 5.25 mM CaCl_2_ for 30 min at 55°C. DNA was purified by phenol-chloroform extraction followed by ethanol precipitation and resuspended in 50 µL nuclease free water.

### ChIP-seq library preparation and sequencing

ChIP-seq libraries were prepared using NEBNext Ultra II DNA library prep kit for Illumina (New England Biolabs) and NEBNext Multiplex Oligos for Illumina (New England Biolabs) according to the manufacturer’s instructions. 5-10 ng ChIP-DNA were used as input for library prep. Sequencing was performed on an Illumina NextSeq 550 instrument in readmode SR75 by the Next Generation Sequencing facility at Vienna BioCenter Core Facilities (VBCF).

### qPCR

qPCR analysis of input and ChIP-DNA was performed on a BioRad CFX Touch cycler using Takyon No Rox SYBR MasterMix dTTP Blue (Eurogentec). qPCR primer sequences are indicated in Supplementary Table 4. Input Cq values were adjusted to 100%, % input values were calculated as follows:

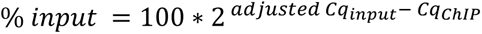

Data are presented as mean ± standard deviation of four replicates. P-value was calculated using a one-tailed, two-sample equal variance t-test. P-values smaller than 0.05 were considered statistically significant, level of significance is indicated with asterisks (‘*’ for p < 0.05; ‘**’ for p < 0.01; ‘***’ for p < 0.001; ‘****’ for p < 0.0001)

### ChIP-seq analysis

ChIP-seq data were mapped to the hg38 version of the human genome using Bowtie2^61^ with the following parameters: bowtie2 -k 1. Human ChIP-seq samples were normalized using mouse chromatin spike-in and mapped to the mouse (mm10) genome, using Bowtie2, as implemented in PigX pipeline^62^. The scaling factor was obtained by dividing the total number of uniquely mapped reads to the human genome, with the number of uniquely mapping reads to the mouse genome. The genomics tracks were then constructed by extending the reads to 200 bp into 3’ direction, calculating the coverage vector, and scaling using the aforementioned scaling factor.

### Data availability

The data that support this study are available from the corresponding author upon reasonable request. The source data are provided with this paper. The atomic coordinates have been deposited in the Protein Data Bank under accession codes: 7Z27 for RBM15 SPOC and 7Z1K for SHARP SPOC:1xS5P CTD. The sequencing data generated in this study have been deposited in ArrayExpress under accession code: E-MTAB-11506 (ChIP-seq). The mass spectrometry proteomics data have been deposited to the ProteomeXchange Consortium via the PRIDE partner repository^63^. The processed mass spectrometry data are provided in Supplementary Datasets 1 and 2. Atomic coordinates used in this study are available in the Protein Data Bank under accession codes 2RT5, 1OW1, 6QV2, 6IC8 and 5KXF and in the Alpha Fold Protein Structure Database under accession codes Q9BTC0, Q6ZMY3 and Q96T37.

## Acknowledgements

We thank Anton Meinhart and Renato Arnese for RBM15 SPOC X-ray data collection; Martin Puchinger for help with labeling CTD peptides and advice regarding fluorescence anisotropy data analysis; Felix Fischer, Kathrin Nagl and Alexander Athanasiadis for help with purifying SPOC domains and FA assays; Dirk Eick for sharing ZNF768 antibody; the VBCF Next Generation Sequencing facility for sequencing; the Max Perutz Labs Mass Spectrometry Facility, Markus Hartl, Dorothea Anrather and Natascha Hartl for mass spectrometry sample processing, data acquisition and analysis; the staff of the X-ray beamlines at ESRF Grenoble and DSL Didcot for their support. L.A. was a member of the Integrative Structural Biology PhD program funded by the Austrian Science Fund (W1258 “DK: Integrative Structural Biology”) from 2018-2021. K.D.C. research was supported by the Wellcome Trust Collaborative Award (201543/Z/16); COST action BM1405 - Non-globular proteins - from sequence to structure, function and application in molecular physiopathology (NGP-NET); WWTF (Vienna Science and Technology Fund) Chemical Biology project LS17-008; Christian Doppler Laboratory for High-Content Structural Biology and Biotechnology; the Austrian-Slovak Interreg Project B301 StruBioMol, University of Vienna Research Platform Comammox and by the University of Vienna. B.Z. is supported by the Austrian Science Fund (P30550). This work was funded by the Austrian Science Fund (P31546 and W1258 “DK: Integrative Structural Biology” to D.S.)

## Author contributions

L.A. purified SPOC domains, performed and analysed fluorescence anisotropy assays, generated endogenously tagged cell lines, performed ChIP-seq experiments, performed and analysed ChIP-qPCR experiments and wrote the manuscript. I.G. solved the SPOC structures. J.B. performed and analysed co-immunoprecipitation assays and analysed mass spectrometry data. V.F. designed and performed the analysis of ChIP-seq data. A.P. performed HADDOCK modelling experiments. A.N. and A.W. purified SPOC domains and performed fluorescence anisotropy assays. B.Z. supervised modelling experiments. A.A. designed and supervised sequencing data analysis. K.D.C. supervised X-ray analysis. D.S. conceived the study, performed, supervised, analysed experiments and wrote the manuscript.

---

**Supplementary Table 1.** X-ray data collection and refinement

|  | **SHARP SPOC:1xS5P CTD** | **RBM15 SPOC** |
| --- | --- | --- |
| Source | MASSIF1 (ESRF, Grenoble, France) | I04 (DLS, Didcot, UK) |
| Wavelength (Å) | 0.9660 | 0.9795 |
| Resolution (Å) | 36.58-1.55<br>(1.58-1.55) | 57.45-1.45<br>(1.47-1.45) |
| Space group | $P2_12_12_1$ | $P2_12_12_1$ |
| Unit cell (Å, °) | 43.33 60.52 68.27 | 43.37 58.97 68.24 |
| Molecules (a.u.) | 1 | 2 |
| Unique reflections | 26393 (1257) | 62616 (4578) |
| Completeness (%) | 98.8 (97.9) | 100 (100) |
| $R_{\text{merge}}^b$ | 0.072 (1.95) | 0.112 (2.08) |
| $R_{\text{meas}}^c$ | 0.083 (2.25) | 0.116 (2.12) |
| CC(1/2) | 0.998 (0.203) | 0.999 (0.571) |
| Multiplicity | 4.3 (4.2) | 3.5 (3.6) |
| $I/\text{sig}(I)$ | 11.3 (1.1) | 12.9 (10.6) |
| $B_{\text{Wilson}} (\text{\AA}^2)$ | 20.4 | 15.7 |
| $R_{\text{work}}^e/R_{\text{free}}^f$ (%) | 21.5/24.7 | 20/21.9 |
| r.m.s.d. bonds (Å) | 0.004 | 0.006 |
| r.m.s.d. angles (°) | 0.731 | 0.872 |

<sup>a</sup> Values in parentheses are for the highest resolution shell.
where $\bar{I}_{(hkl)}$ is the mean intensity of multiple $I_{i(hkl)}$ observations of the symmetry-related reflections, N is the redundancy
<sup>f</sup> $R_{\text{free}}$ is the cross-validation $R_{\text{factor}}$ computed for the randomly chosen test set of reflections (5 %) which are omitted in the refinement process.

**Supplementary Table 2.**
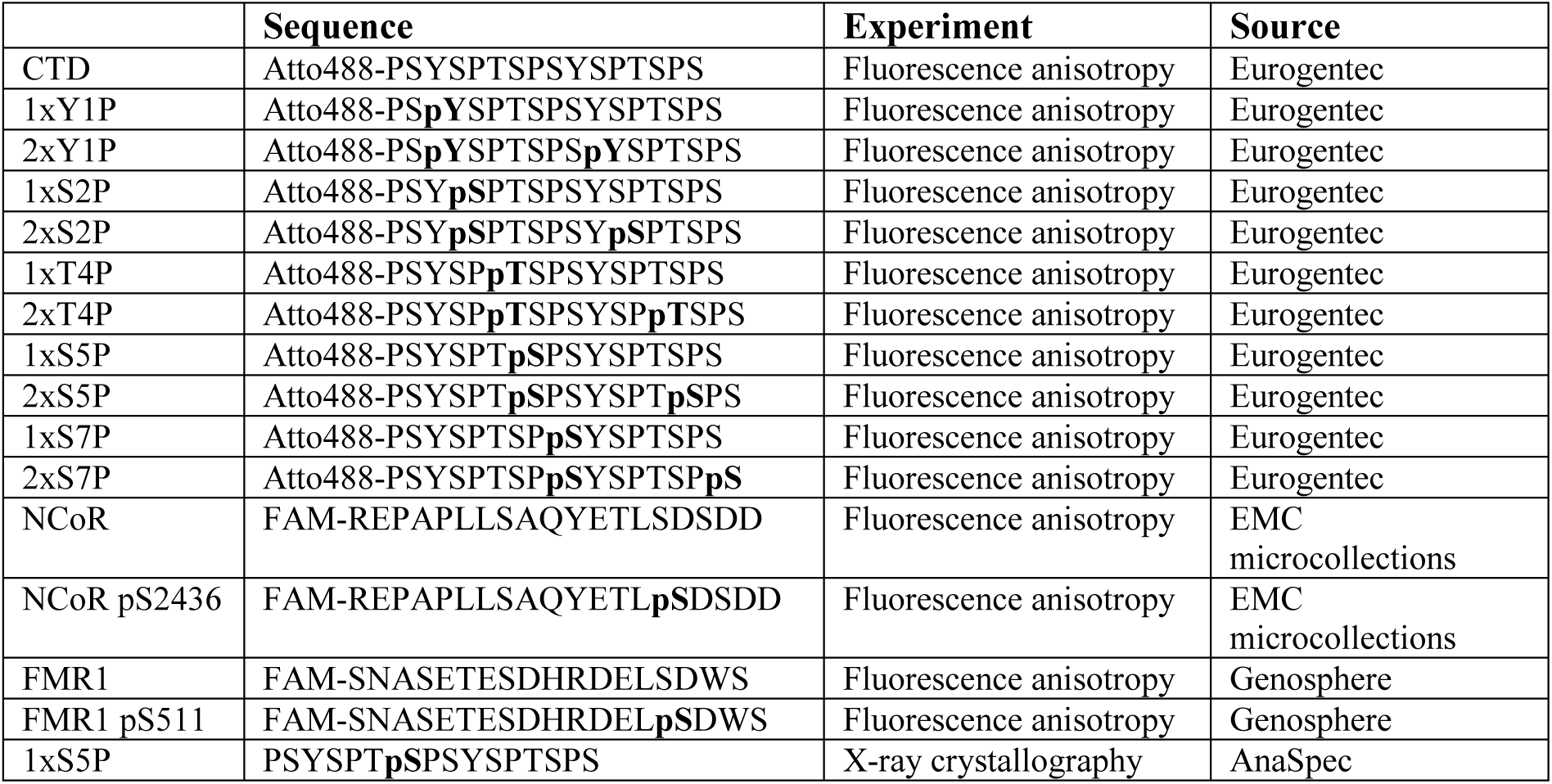
Peptides

**Supplementary Table 3.**
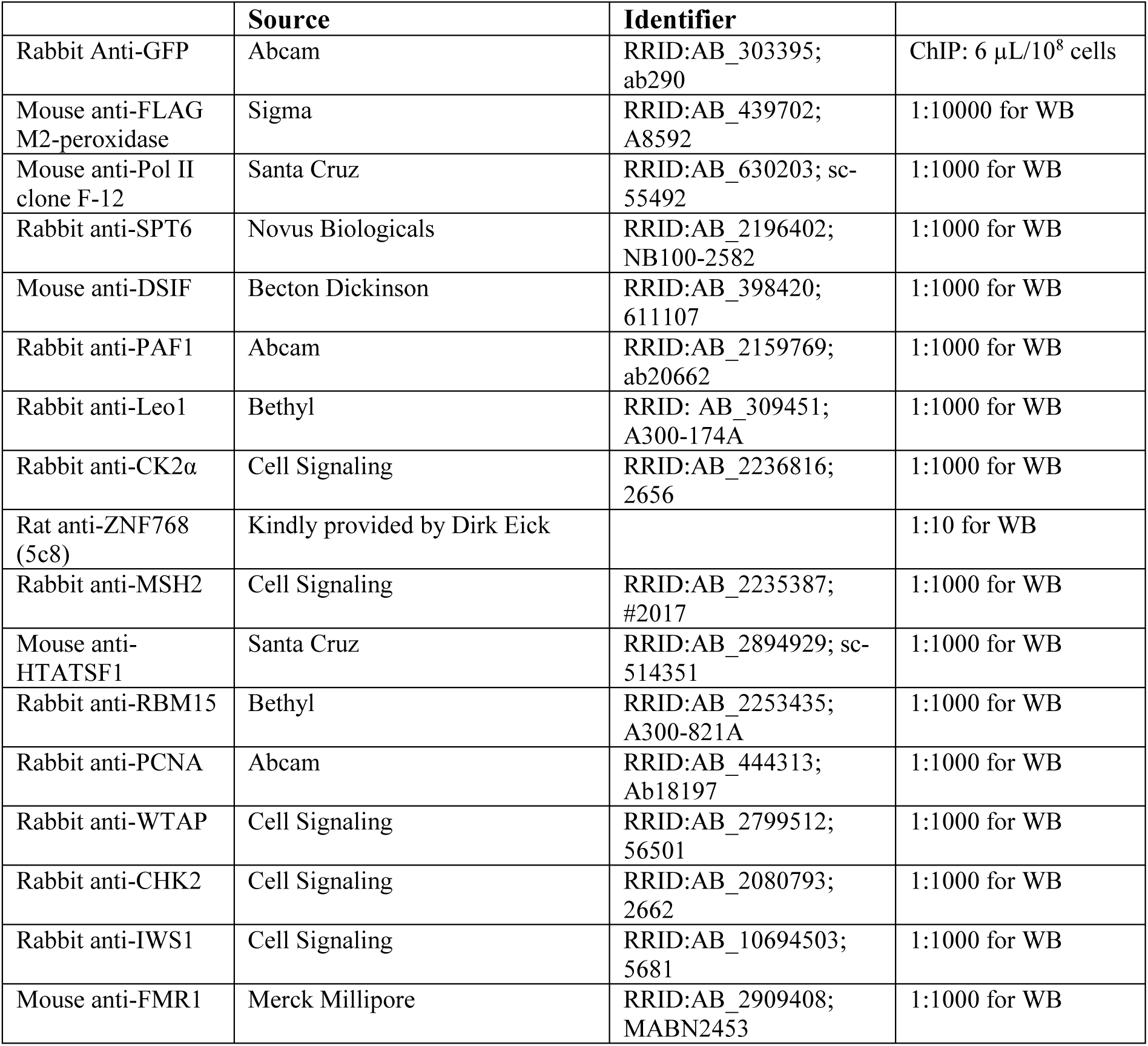
Antibodies

**Supplementary Table 4.**
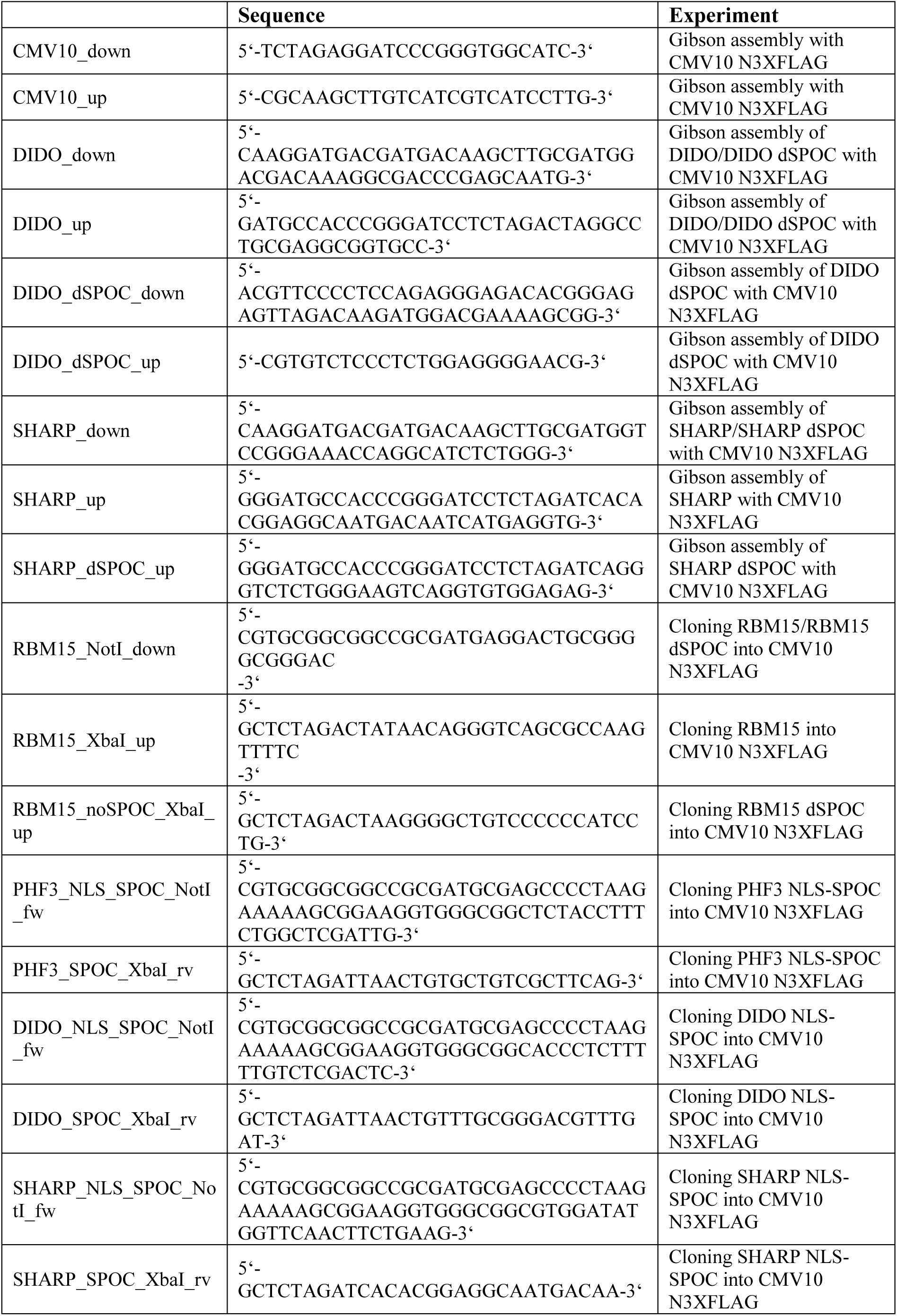
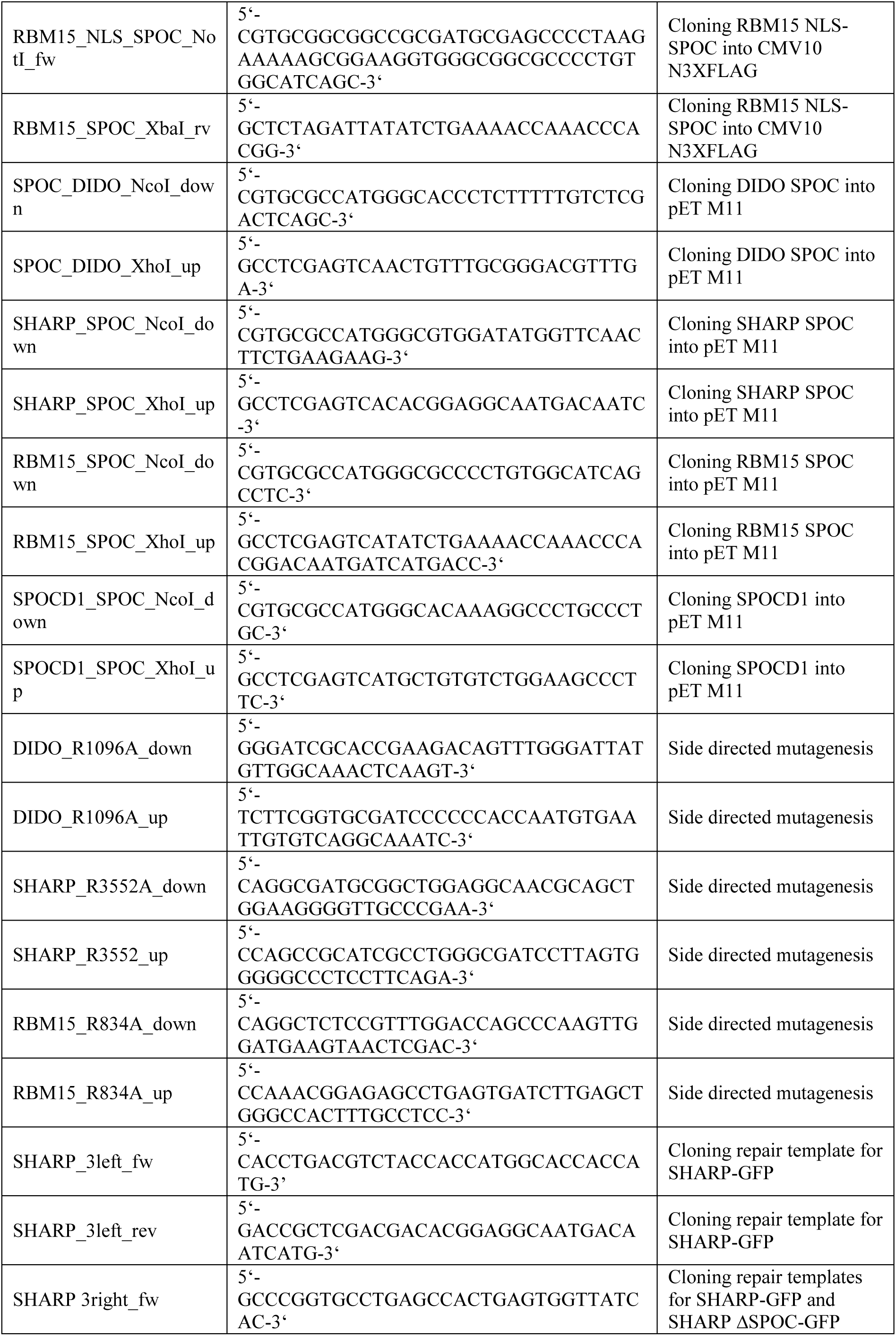
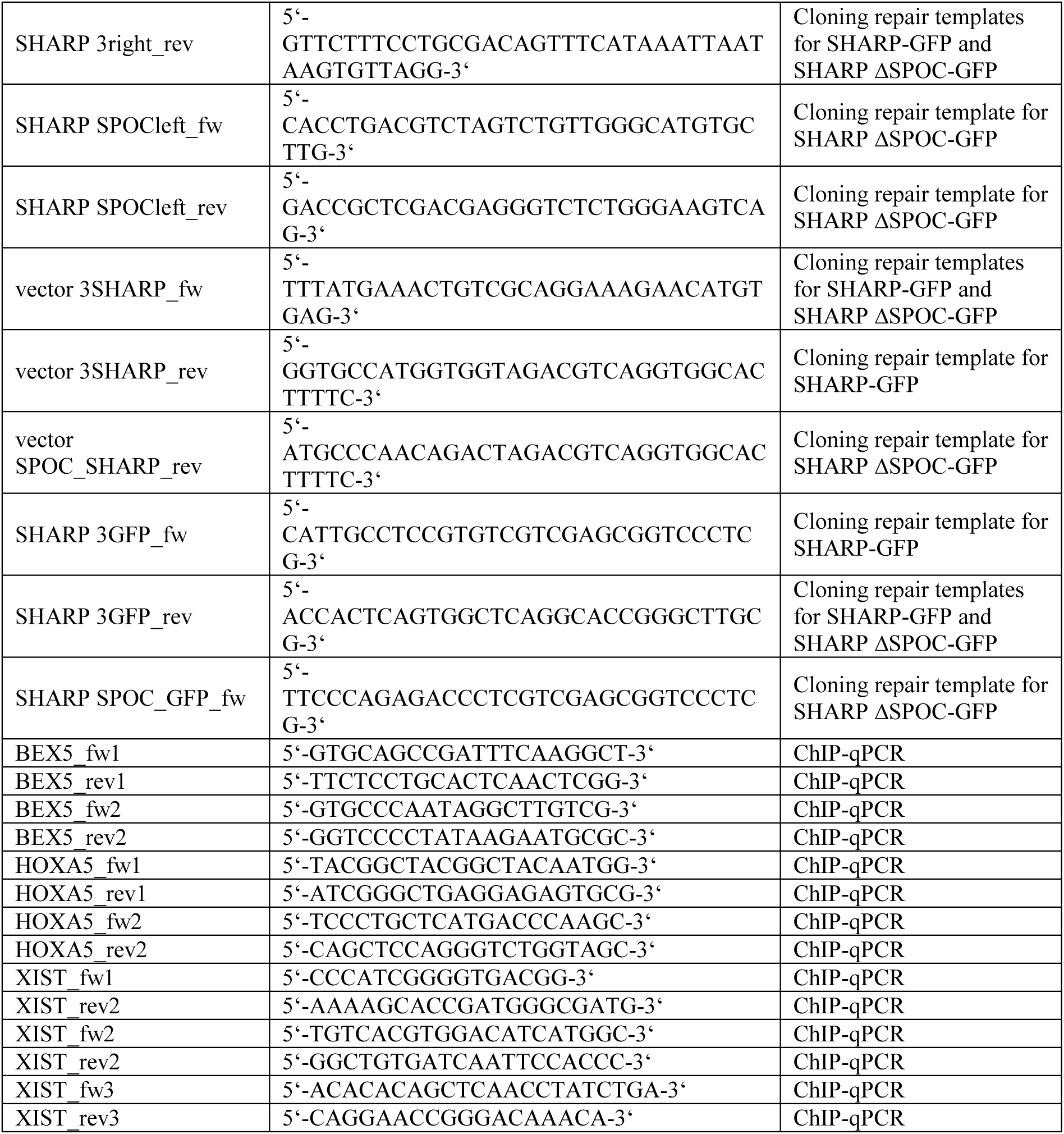
Oligonucleotides

**Supplementary Fig. 1:**
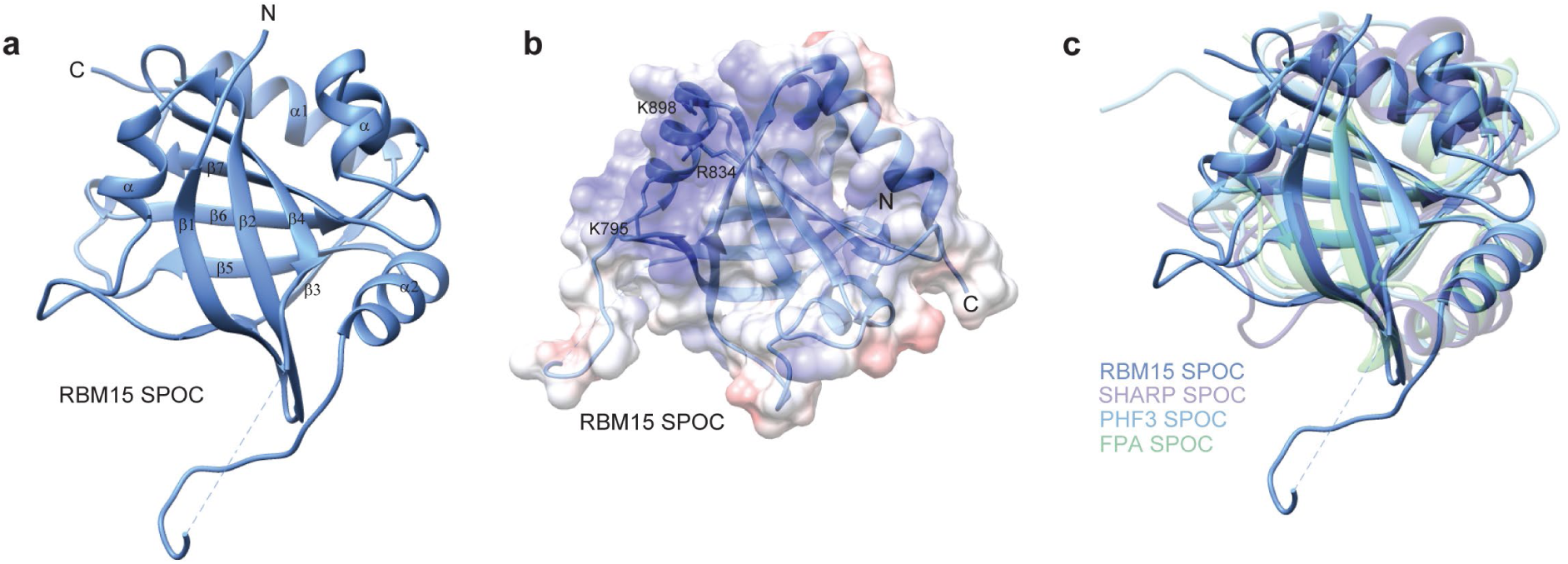
Crystal structure of the RBM15 SPOC domain. **a** Ribbon model of the crystal structure of RBM15 SPOC. **b** Overlay of RBM15 SPOC ribbon model and electrostatic surface potential. Conserved lysine and arginine residues of the basic patch are indicated. Electrostatic surface potential was calculated using the Coulombic Surface Coloring tool in UCSF Chimera and is depicted ranging from -10 (red) to +10 (blue) kcal/(mol*e). **c** Overlay of SPOC structures from RBM15 (7Z27), SHARP (2RT5), PHF3 (6Q2V) and FPA (5KXF) showed an average RMSD of 1.075 Å over 99 pruned Cα atoms between RBM15 and SHARP SPOC (6.645 Å over all 159 pairs), an average RMSD of 1.075 Å over 72 pruned Cα atoms between RBM15 and PHF3 SPOC (6.387 Å over all 145 pairs) and an average RMSD of 0.913 Å over 53 pruned Cα atoms between RBM15 and FPA SPOC (5.531 Å over all 118 pairs). The alignment was generated using the Matchmaker tool in UCSF Chimera.

**Supplementary Fig. 2:**
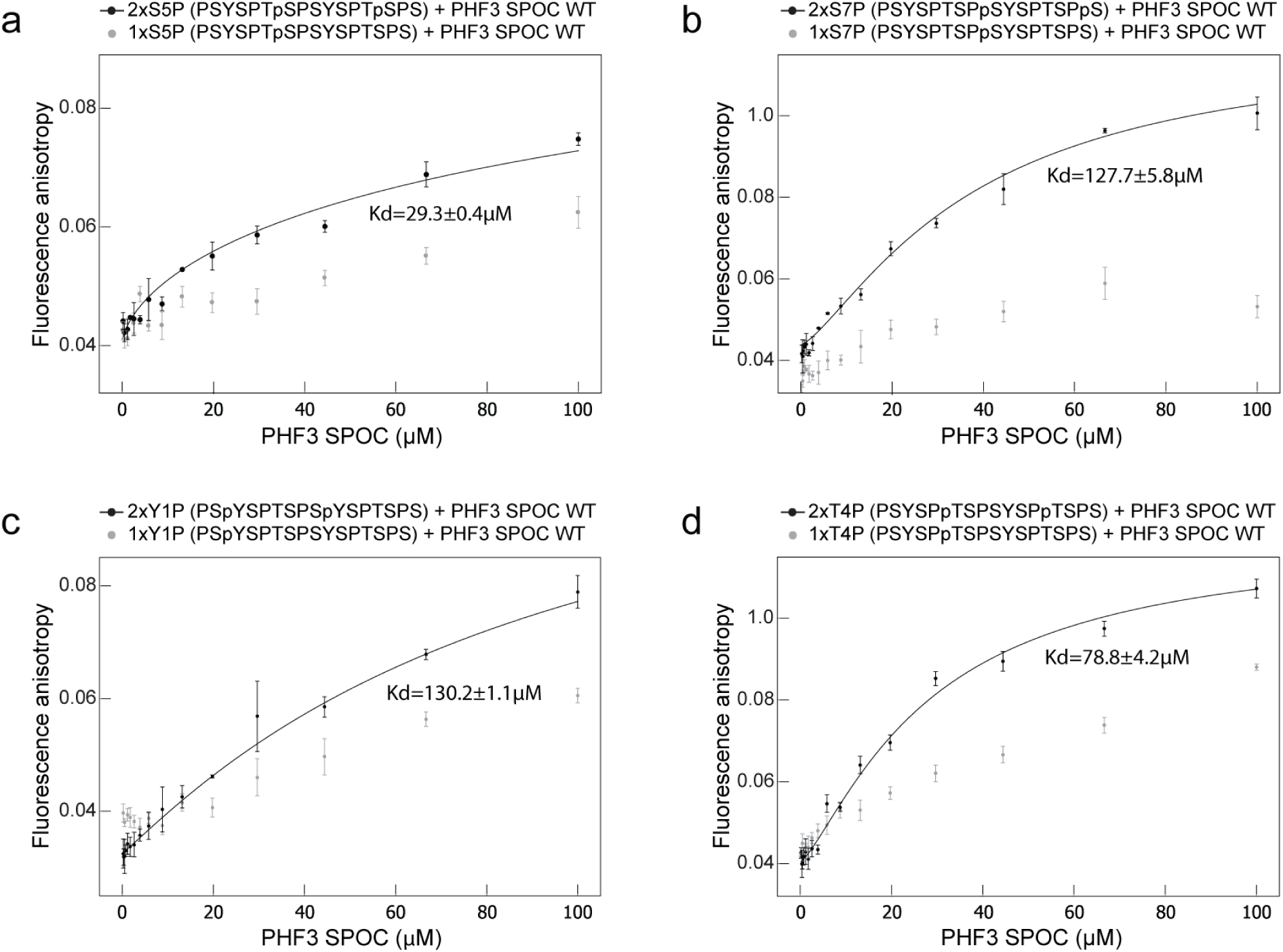
PHF3 SPOC-CTD binding assays. Fluorescence anisotropy measurements of PHF3 SPOC with CTD peptides phosphorylated on **a** serine-5, **b** serine-7, **c** tyrosine-1 and **d** threonine-4. Fluorescence anisotropy is plotted as a function of protein concentration. Data points and error bars represent the mean ± standard deviation from 3 independent experiments.

**Supplementary Fig. 3:**
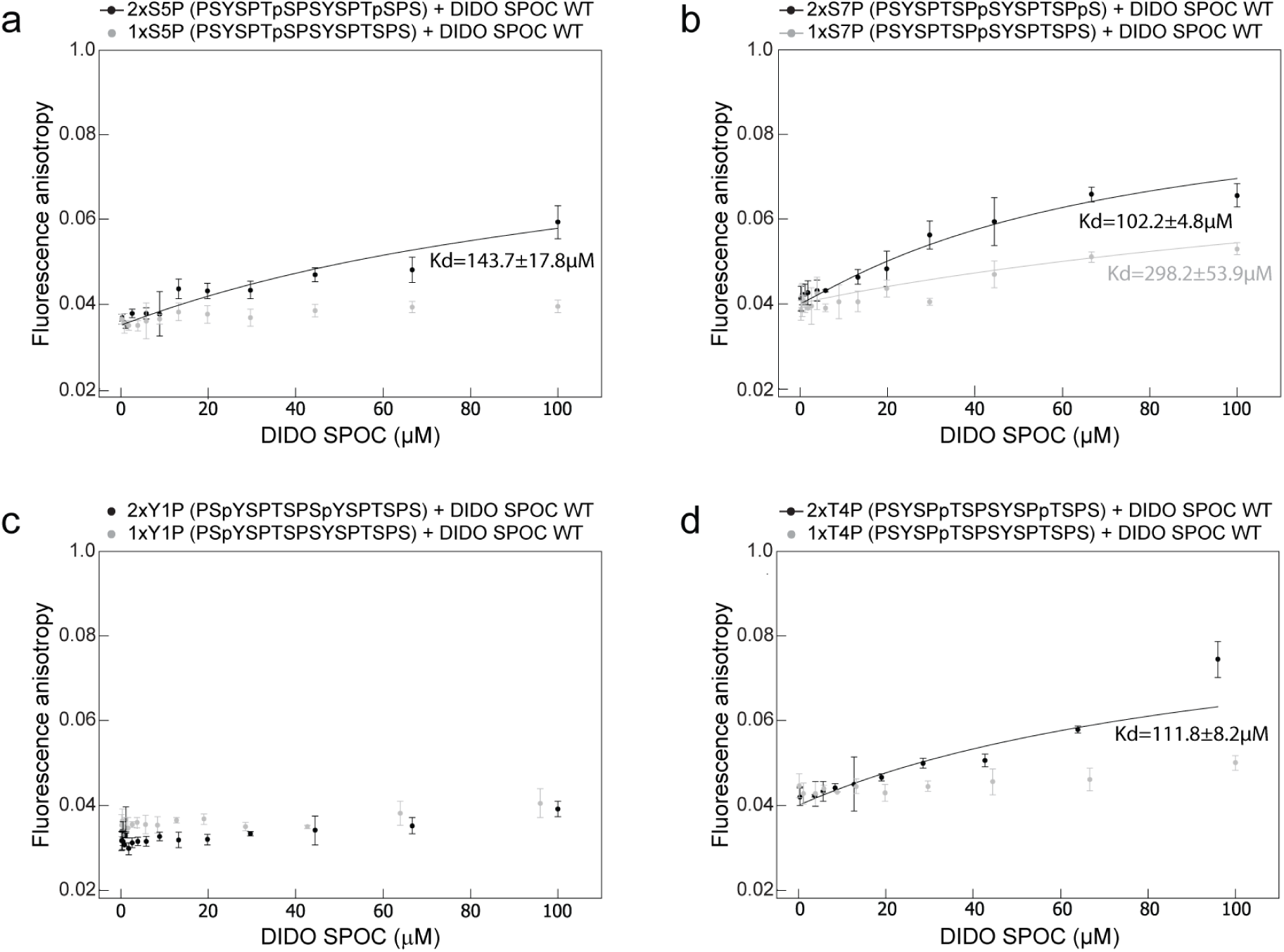
DIDO SPOC-CTD binding assays. Fluorescence anisotropy measurements of DIDO SPOC with CTD peptides phosphorylated on **a** serine-5, **b** serine-7, **c** tyrosine-1 and **d** threonine-4. Fluorescence anisotropy is plotted as a function of protein concentration. Data points and error bars represent the mean ± standard deviation from 3 independent experiments.

**Supplementary Fig. 4:**
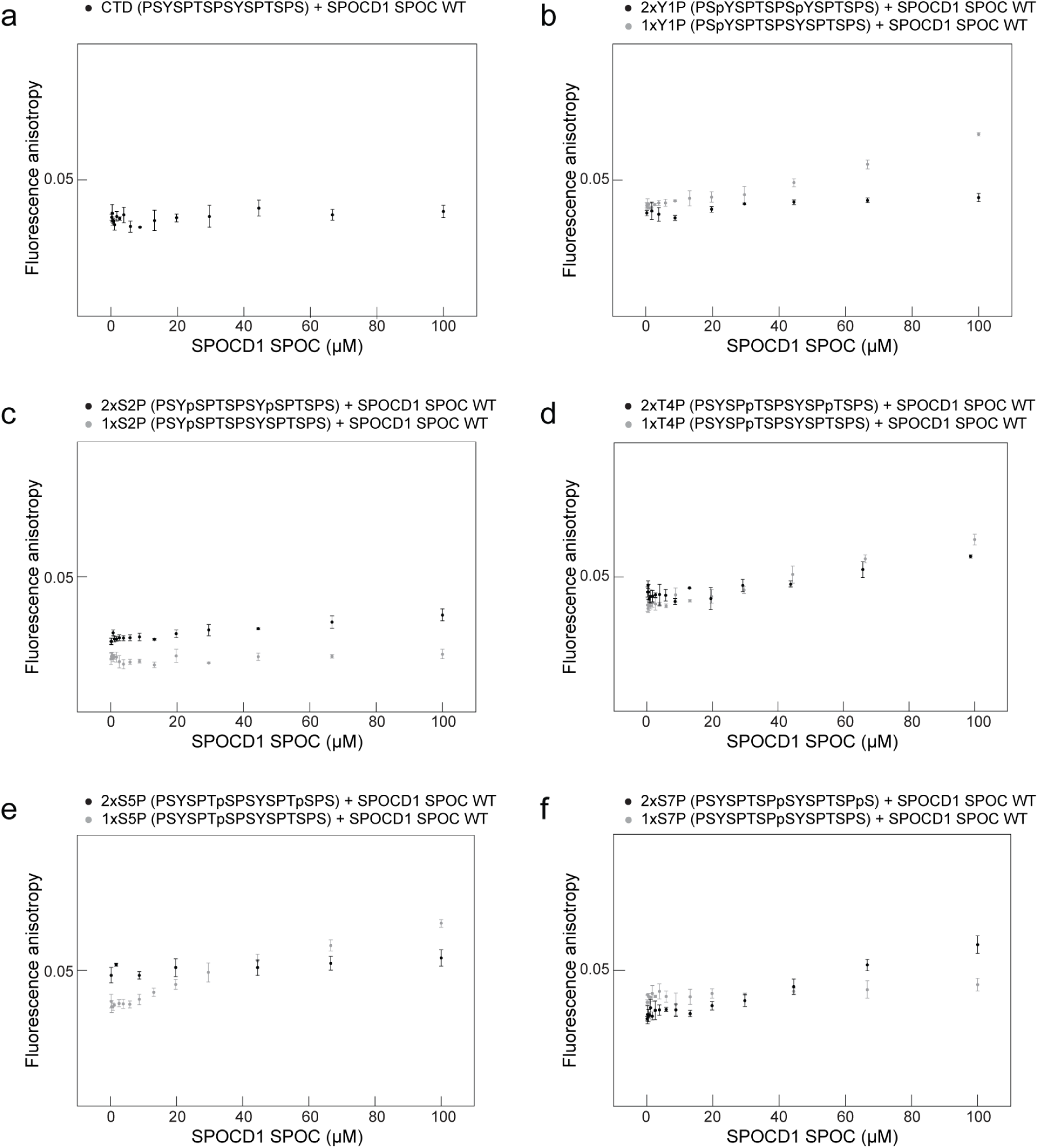
SPOCD1 SPOC-CTD binding assays. Fluorescence anisotropy measurements of SPOCD1 SPOC with **a** unphosphorylated CTD peptide or CTD peptides phosphorylated on **b** tyrosine-1, **c** serine-2, **d** threonine-4, **e** serine-5 and **f** serine-7. Fluorescence anisotropy is plotted as a function of protein concentration. Data points and error bars represent the mean ± standard deviation from 3 independent experiments.

**Supplementary Fig. 5:**
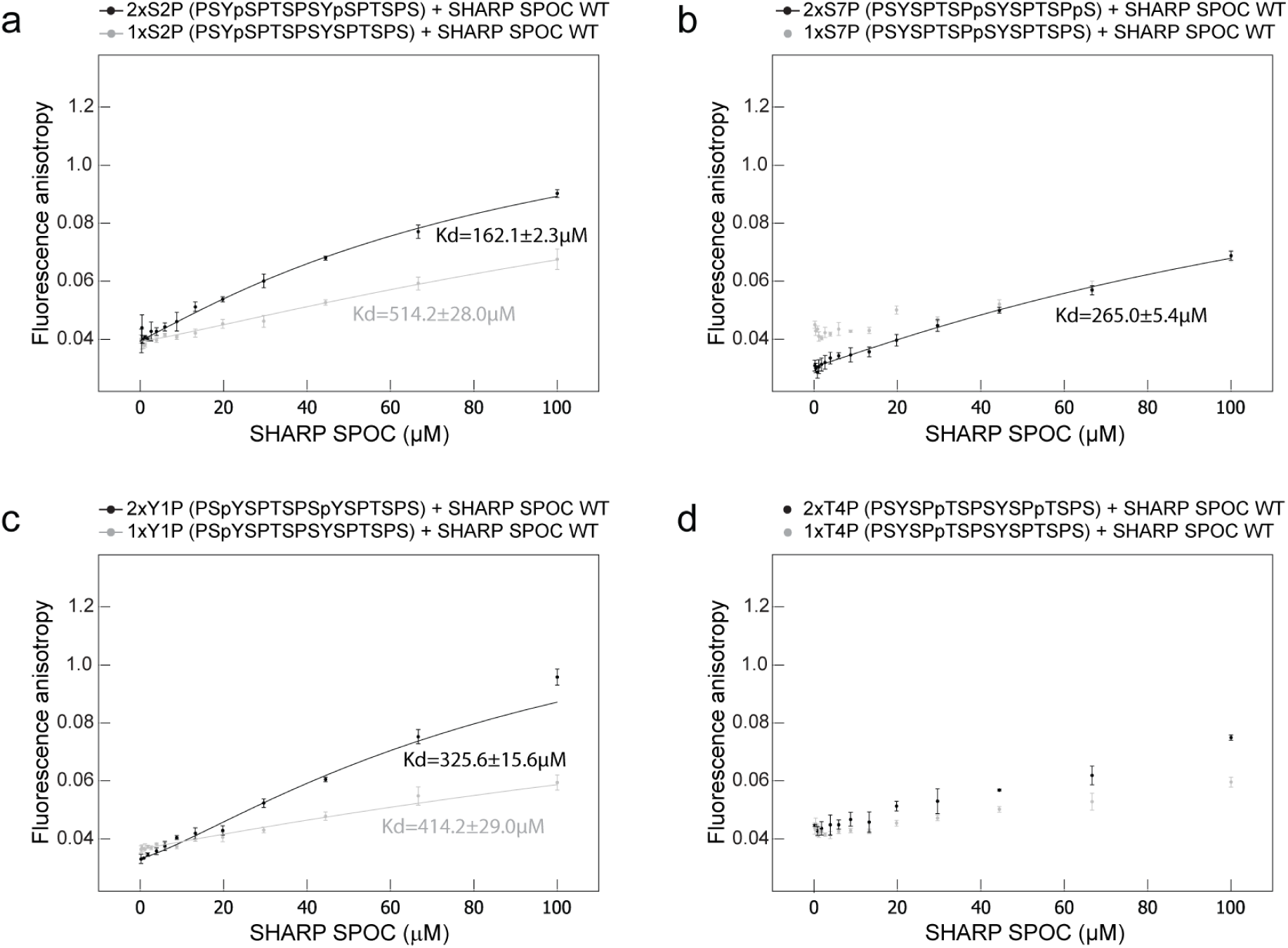
SHARP SPOC-CTD binding assays. Fluorescence anisotropy measurements of SHARP SPOC with CTD peptides phosphorylated on **a** serine-2, **b** serine-7, **c** tyrosine-1 and **d** threonine-4. Fluorescence anisotropy is plotted as a function of protein concentration. Data points and error bars represent the mean ± standard deviation from 3 independent experiments.

**Supplementary Fig. 6:**
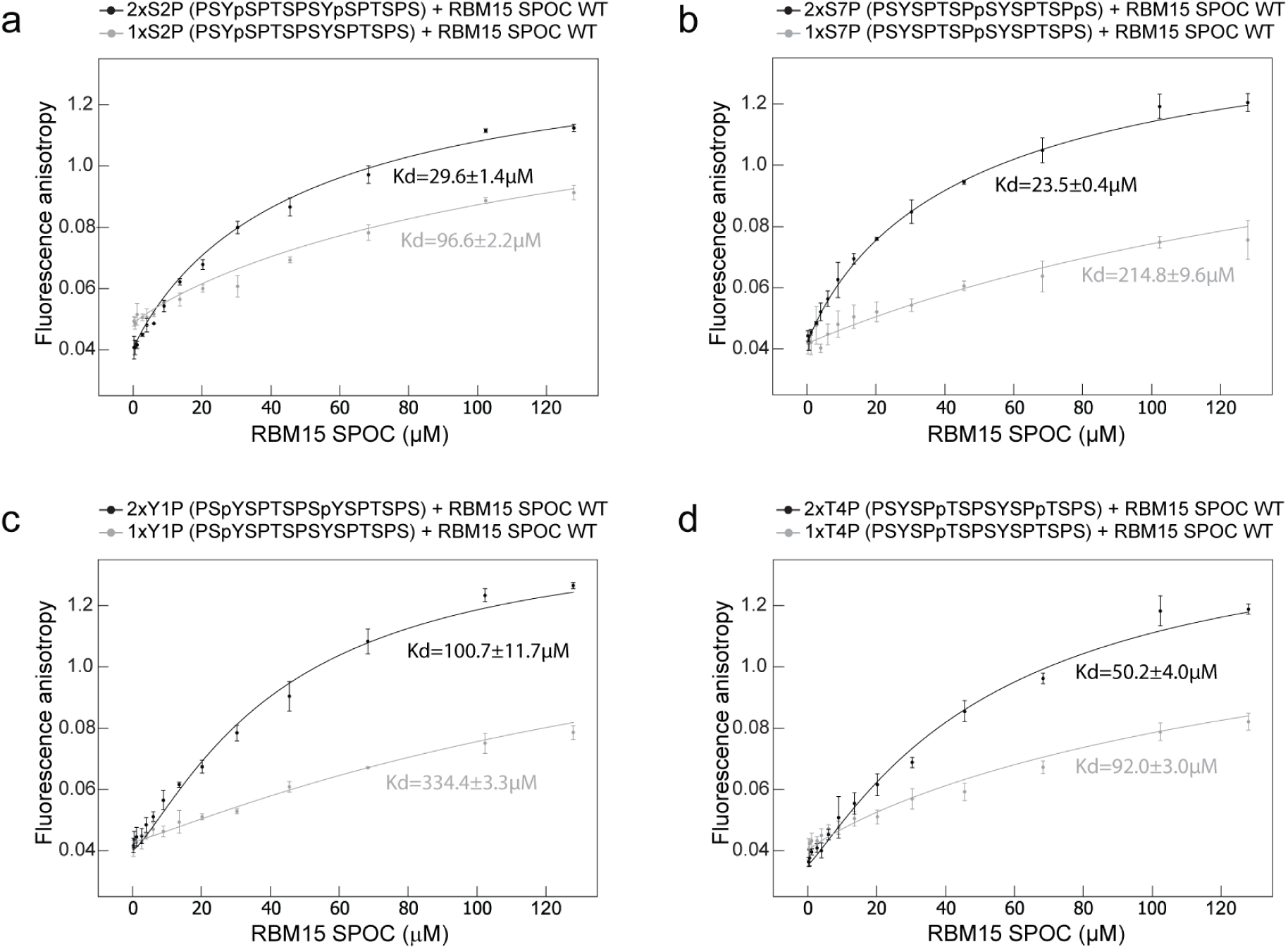
RBM15 SPOC-CTD binding assays. Fluorescence anisotropy measurements of RBM15 SPOC with CTD peptides phosphorylated on **a** serine-2, **b** serine-7, **c** tyrosine-1 and **d** threonine-4. Fluorescence anisotropy is plotted as a function of protein concentration. Data points and error bars represent the mean ± standard deviation from 3 independent experiments.

**Supplementary Fig. 7:**
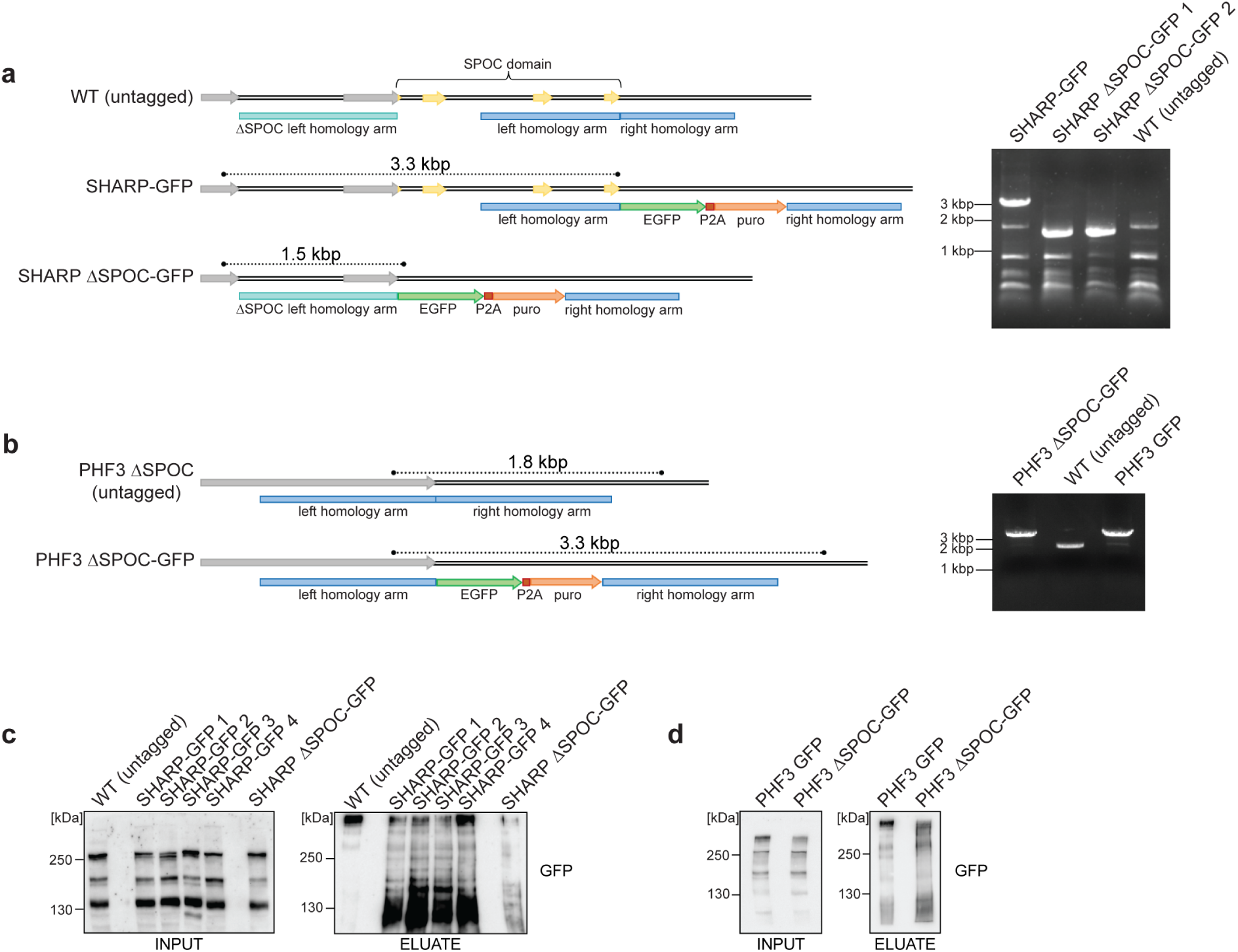
Generation of endogenously tagged SHARP-GFP and PHF3-GFP cell lines. **a** CRISPR/Cas9 strategy and PCR validation of endogenous tagging of SHARP with GFP at the C-terminus (SHARP-GFP) and deletion of the SPOC domain with simultaneous C-terminal GFP-tagging (SHARP ΔSPOC-GFP). The experiment was performed once. Genotyping is shown for two individual SHARP ΔSPOC-GFP clones. **b** CRISPR/Cas9 strategy and PCR validation of endogenous tagging of PHF3 ΔSPOC with GFP at the C-terminus (PHF3 ΔSPOC-GFP). The experiment was performed once. The PHF3 ΔSPOC cell line had been generated and validated before. **c** GFP-IP of endogenous SHARP-GFP and SHARP ΔSPOC-GFP. The experiment was performed once. Four individual clones are shown for SHARP-GFP. **d** GFP-IP of endogenous PHF3-GFP and PHF3 ΔSPOC-GFP. The experiment was performed once.

**Supplementary Fig. 8:**
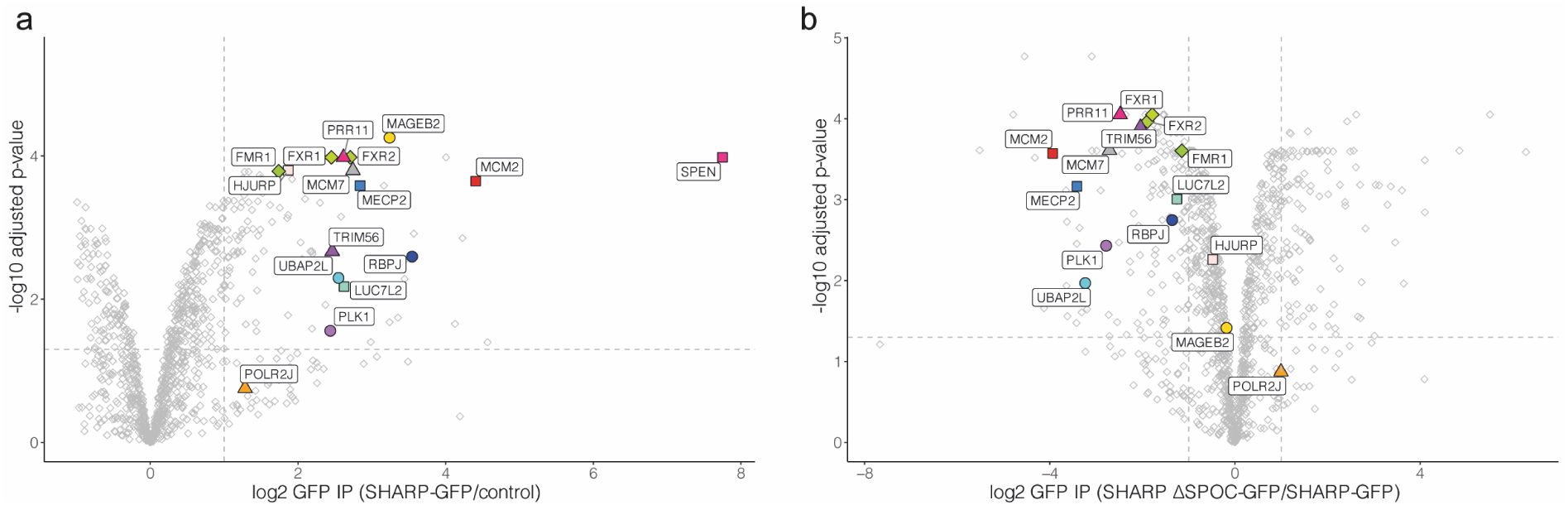
Interactome of endogenous SHARP-GFP. Volcano plots of SHARP-GFP interactors identified by mass spectrometry **a** compared to an untagged control cell line and **b** compared to SHARP ΔSPOC-GFP. The experiments were performed in three replicates.

## References

1. Sánchez-Pulido, L., Rojas, A. M., van Wely, K. H., Martinez-A, C. & Valencia, A. SPOC: A widely distributed domain associated with cancer, apoptosis and transcription. BMC Bioinformatics 5, 6–11 (2004).

2. Ariyoshi, M. & Schwabe, J. W. R. A conserved structural motif reveals the essential transcriptional repression function of spen proteins and their role in developmental signaling. Genes and Development 17, 1909–1920 (2003).

3. Mikami, S. et al. Structural insights into the recruitment of SMRT by the corepressor SHARP under phosphorylative regulation. Structure 22, 35–46 (2014).

4. Zhang, Y., Rataj, K., Simpson, G. G. & Tong, L. Crystal Structure of the SPOC Domain of the Arabidopsis Flowering Regulator FPA. PLOS ONE 1–13 (2016) doi:10.1371/journal.pone.0160694.

5. Appel, L.-M. et al. PHF3 regulates neuronal gene expression through the Pol II CTD reader domain SPOC. Nature Communications 12, 6078 (2021).

6. Zoch, A. et al. SPOCD1 is an essential executor of piRNA-directed de novo DNA methylation. Nature 584, 635–639 (2020).

7. Oswald, F. et al. RBP-Jkappa/SHARP recruits CtIP/CtBP corepressors to silence Notch target genes. Molecular and cellular biology 25, 10379–10390 (2005).

8. Oswald, F. et al. SHARP is a novel component of the Notch/RBP-Jκ signalling pathway. EMBO Journal 21, 5417–5426 (2002).

9. Villares, R. et al. Dido mutations trigger perinatal death and generate brain abnormalities and behavioral alterations in surviving adult mice. Proceedings of the National Academy of Sciences of the United States of America 112, 4803–4808 (2015).

10. Ma, X. et al. Rbm15 modulates Notch-induced transcriptional activation and affects myeloid differentiation. Molecular and cellular biology 27, 3056–3064 (2007).

11. Oswald, F. et al. A phospho-dependent mechanism involving NCoR and KMT2D controls a permissive chromatin state at Notch target genes. Nucleic Acids Research 44, 4703–4720 (2016).

12. Dossin, F. et al. SPEN integrates transcriptional and epigenetic control of X-inactivation. Nature 578, 455–460 (2020).

13. Hiriart, E. et al. Interaction of the Epstein-Barr Virus mRNA Export Factor EB2 with Human Spen Proteins SHARP, OTT1, and a Novel Member of the Family, OTT3, Links Spen Proteins with Splicing Regulation and mRNA Export *. Journal of Biological Chemistry 280, 36935–36945 (2005).

14. Uranishi, H. et al. The RNA-binding motif protein 15B (RBM15B/OTT3) acts as cofactor of the nuclear export receptor NXF1. The Journal of biological chemistry 284, 26106–26116 (2009).

15. Zhang, L. et al. Cross-talk between PRMT1-mediated methylation and ubiquitylation on RBM15 controls RNA splicing. eLife 4, e07938 (2015).

16. Knuckles, P. et al. Zc3h13/Flacc is required for adenosine methylation by bridging the mRNA-binding factor Rbm15/Spenito to the m(6)A machinery component Wtap/Fl(2)d. Genes & development 32, 415–429 (2018).

17. Lee, J.-H. & Skalnik, D. G. Rbm15-Mkl1 Interacts with the Setd1b Histone H3-Lys4 Methyltransferase via a SPOC Domain That Is Required for Cytokine-Independent Proliferation. PLOS ONE 7, e42965- (2012).

18. Harlen, K. M. & Churchman, L. S. The code and beyond : transcription regulation by the RNA polymerase II carboxy-terminal domain. Nature Reviews, Molecular Cell Biology 18, 263–273 (2017).

19. Eick, D. & Geyer, M. The RNA Polymerase II Carboxy-Terminal Domain (CTD) Code. Chemical Reviews 113, 8456–8490 (2013).

20. Ghosh, A., Shuman, S. & Lima, C. D. Structural insights to how mammalian capping enzyme reads the CTD code. Molecular cell 43, 299–310 (2011).

21. Meinhart, A. & Cramer, P. Recognition of RNA polymerase II carboxy-terminal domain by 3′-RNA-processing factors. Nature 430, 223–226 (2004).

22. Fütterer, A. et al. DIDO as a Switchboard that Regulates Self-Renewal and Differentiation in Embryonic Stem Cells. Stem cell reports 8, 1062–1075 (2017).

23. Fütterer, A. et al. Ablation of Dido3 compromises lineage commitment of stem cells in vitro and during early embryonic development. Cell death and differentiation 19, 132– 143 (2012).

24. Mora Gallardo, C., et al. Dido3-dependent SFPQ recruitment maintains efficiency in mammalian alternative splicing. Nucleic acids research 47, 5381–5394 (2019).

25. Mora Gallardo, C., Sánchez de Diego, A., Martínez-A, C. & van Wely, K. H. M. Interplay between splicing and transcriptional pausing exerts genome-wide control over alternative polyadenylation. Transcription 12, 55–71 (2021).

26. Varadi, M. et al. AlphaFold Protein Structure Database: massively expanding the structural coverage of protein-sequence space with high-accuracy models. Nucleic acids research 50, D439–D444 (2022).

27. Jumper, J. et al. Highly accurate protein structure prediction with AlphaFold. Nature 596, 583–589 (2021).

28. van Zundert, G. C. P. et al. The HADDOCK2.2 Web Server: User-Friendly Integrative Modeling of Biomolecular Complexes. Journal of Molecular Biology 428, 720–725 (2016).

29. Honorato, R. v, et al. Structural Biology in the Clouds: The WeNMR-EOSC Ecosystem. Frontiers in molecular biosciences 8, 729513 (2021).

30. Rohrmoser, M. et al. MIR sequences recruit zinc finger protein ZNF768 to expressed genes. Nucleic Acids Research 47, 700–715 (2019).

31. Meggio, F. & Pinna, L. A. One-thousand-and-one substrates of protein kinase CK2? The FASEB Journal 17, 349–368 (2003).

32. Zhou, Y., Gross, W., Hong, S. H. & Privalsky, M. L. The SMRT corepressor is a target of phosphorylation by protein kinase CK2 (casein kinase II). Molecular and cellular biochemistry 220, 1–13 (2001).

33. Loerch, S. et al. The pre-mRNA splicing and transcription factor Tat-SF1 is a functional partner of the spliceosome SF3b1 subunit via a U2AF homology motif interface. The Journal of biological chemistry 294, 2892–2902 (2019).

34. Edens, B. M. et al. FMRP Modulates Neural Differentiation through m(6)A-Dependent mRNA Nuclear Export. Cell reports 28, 845–854.e5 (2019).

35. Majumder, M., Johnson, R. H. & Palanisamy, V. Fragile X-related protein family: a double-edged sword in neurodevelopmental disorders and cancer. Critical reviews in biochemistry and molecular biology 55, 409–424 (2020).

36. Hornbeck, P. v, et al. PhosphoSitePlus, 2014: mutations, PTMs and recalibrations. Nucleic acids research 43, D512–D520 (2015).

37. Kinkelin, K. et al. Structures of RNA polymerase II complexes with Bye1, a chromatin-binding PHF3/DIDO homologue. Proceedings of the National Academy of Sciences 110, 15277 (2013).

38. Cheung, A. C. M. & Cramer, P. Structural basis of RNA polymerase II backtracking, arrest and reactivation. Nature 471, 249–253 (2011).

39. Gatchalian, J. et al. Dido3 PHD modulates cell differentiation and division. Cell reports 4, 148–158 (2013).

40. Tencer, A. H. et al. A Unique pH-Dependent Recognition of Methylated Histone H3K4 by PPS and DIDO. Structure (London, England : 1993) 25, 1530–1539.e3 (2017).

41. Chu, C. et al. Systematic Discovery of Xist RNA Binding Proteins. Cell 161, 404–416 (2015).

42. Monfort, A. et al. Identification of Spen as a Crucial Factor for Xist Function through Forward Genetic Screening in Haploid Embryonic Stem Cells. Cell reports 12, 554– 561 (2015).

43. Lu, Z. et al. Resource RNA Duplex Map in Living Cells Reveals Higher-Order Transcriptome Structure Resource RNA Duplex Map in Living Cells Reveals Higher-Order Transcriptome Structure. Cell 165, 1267–1279 (2016).

44. McHugh, C. A. et al. The Xist lncRNA Interacts Directly With SHARP to Silence Transcription Through HDAC3. Nature 521, 232–236 (2015).

45. Nesterova, T. B. et al. Systematic allelic analysis defines the interplay of key pathways in X chromosome inactivation. Nature Communications 10, 3129 (2019).

46. Chaumeil, J., le Baccon, P., Wutz, A. & Heard, E. A novel role for Xist RNA in the formation of a repressive nuclear compartment into which genes are recruited when silenced. Genes & development 20, 2223–2237 (2006).

47. Bach, S., Shovlin, S., Moriarty, M., Bardoni, B. & Tropea, D. Rett Syndrome and Fragile X Syndrome: Different Etiology With Common Molecular Dysfunctions. Frontiers in cellular neuroscience 15, 764761 (2021).

48. Chen, E. & Joseph, S. Fragile X mental retardation protein: A paradigm for translational control by RNA-binding proteins. Biochimie 114, 147–154 (2015).

49. Shi, H., Wei, J. & He, C. Where, When, and How: Context-Dependent Functions of RNA Methylation Writers, Readers, and Erasers. Molecular cell 74, 640–650 (2019).

50. Patil, D. P. et al. m6A RNA methylation promotes XIST-mediated transcriptional repression. Nature 537, 369–373 (2016).

51. Li, C. et al. FastCloning: a highly simplified, purification-free, sequence- and ligation-independent PCR cloning method. BMC Biotechnology 11, 92 (2011).

52. Kabsch, W. XDS. Acta crystallographica. Section D, Biological crystallography 66, 125–132 (2010).

53. Winn, M. D. et al. Overview of the CCP4 suite and current developments. Acta crystallographica. Section D, Biological crystallography 67, 235–242 (2011).

54. Adams, P. D. et al. PHENIX: a comprehensive Python-based system for macromolecular structure solution. Acta crystallographica. Section D, Biological crystallography 66, 213–221 (2010).

55. Emsley, P. & Cowtan, K. Coot: model-building tools for molecular graphics. Acta Crystallographica Section D Biological Crystallography 60, 2126–2132 (2004).

56. Cong, L. et al. Multiplex genome engineering using CRISPR/Cas systems. Science (New York, N.Y.) 339, 819–823 (2013).

57. Rappsilber, J., Mann, M. & Ishihama, Y. Protocol for micro-purification, enrichment, pre-fractionation and storage of peptides for proteomics using StageTips. Nature Protocols 2, 1896–1906 (2007).

58. Tyanova, S., Temu, T. & Cox, J. The MaxQuant computational platform for mass spectrometry-based shotgun proteomics. Nature Protocols 11, 2301–2319 (2016).

59. R Core Team. R: A language and environment for statistical computing. R Foundation for Statistical Computing, Vienna, Austria. Available online at https://www.R-project.org/. (2018).

60. Ritchie, M. E. et al. limma powers differential expression analyses for RNA-sequencing and microarray studies. Nucleic Acids Research 43, e47–e47 (2015).

61. Langmead, B. & Salzberg, S. L. Fast gapped-read alignment with Bowtie 2. Nature methods 9, 357–359 (2012).

62. Wurmus, R. et al. PiGx: reproducible genomics analysis pipelines with GNU Guix. GigaScience 7, giy123 (2018).

63. Perez-Riverol, Y. et al. The PRIDE database and related tools and resources in 2019: improving support for quantification data. Nucleic acids research 47, D442–D450 (2019).

64. Pettersen, E. F. et al. UCSF Chimera - A visualization system for exploratory research and analysis. Journal of Computational Chemistry 25, 1605–1612 (2004).

